# Reducing reference bias using multiple population reference genomes

**DOI:** 10.1101/2020.03.03.975219

**Authors:** Nae-Chyun Chen, Brad Solomon, Taher Mun, Sheila Iyer, Ben Langmead

## Abstract

Most sequencing data analyses start by aligning sequencing reads to a linear reference genome. But failure to account for genetic variation causes reference bias and confounding of results downstream. Other approaches replace the linear reference with structures like graphs that can include genetic variation, incurring major computational overhead. We propose the “reference flow” alignment method that uses multiple population reference genomes to improve alignment accuracy and reduce reference bias. Compared to the graph aligner vg, reference flow achieves a similar level of accuracy and bias avoidance, but with 14% of the memory footprint and 5.5 times the speed.

## 1 Introduction

Sequencing data analysis often begins with aligning reads to a reference genome, with the reference represented as a linear string of bases. Linear references such as the primary GRCh38 assembly^1^ work naturally with efficient text indexes and sequence alignment algorithms. But linearity leads to *reference bias*: a tendency to miss alignments or report incorrect alignments for reads containing non-reference alleles. This can ultimately lead to confounding of scientific results, especially for analyses concerned with hypervariable regions^2^, allele-specific effects^3–6^, ancient DNA analysis^7,8^ or epigenenomic signals^9^. These problems can be more or less adverse depending on the individual under study; e.g. African-ancestry genomes contain more ALT alleles, and so can be more severely affected by reference bias^10^.

While graph aligners^11–15^ can reduce reference bias, linear aligners still perform better on certain classes of reads^16^ and graph-aligner performance is sensitive to the number of variants considered^17^. Other efforts have focused on elaborating the linear alignment paradigm to address reference bias. Some studies suggest replacing the typical linear reference with a “major-allele” version, with each variant set to its most common allele. This can increase alignment^16–18^ and genotyping accuracy^19^. The major-allele reference is largely compatible with the standard reference (though indels can shift coordinates) and imposes little or no additional computational overhead.

We propose a new strategy called “reference flow” that uses a collection of references chosen so as to cover known genetic variants (Figure 1). We call the method “reference flow” because it selects which reads to align to which genomes based on how well the read aligned previously. In this work, we propose specific reference-flow strategies where the method proceeds in two passes where the first pass aligns reads to the “initial” reference and identifies unaligned reads and reads with ambiguous (low mapping-quality) alignments. The second pass re-aligns these reads to a collection of references that are chosen to span the genetic space. By merging results from both passes, we can achieve higher alignment sensitivity and lower reference bias compared to methods that use a single reference. We implemented methods to align second-pass reads to the set of five genomes corresponding to the “super populations” studied in the 1000 Genomes Project^20^, as well as to the set of 26 genomes corresponding to the more specific 1000 Genomes “populations.” This method (a) can use existing, robust linear aligners like Bowtie 2^21^ or BWA-MEM^22^, (b) requires only a small number of pre-established linear reference genomes, and (c) imposes minimal computational overhead – with no possibility of exponential blowup – relative to linear aligners.

**Figure 1:**
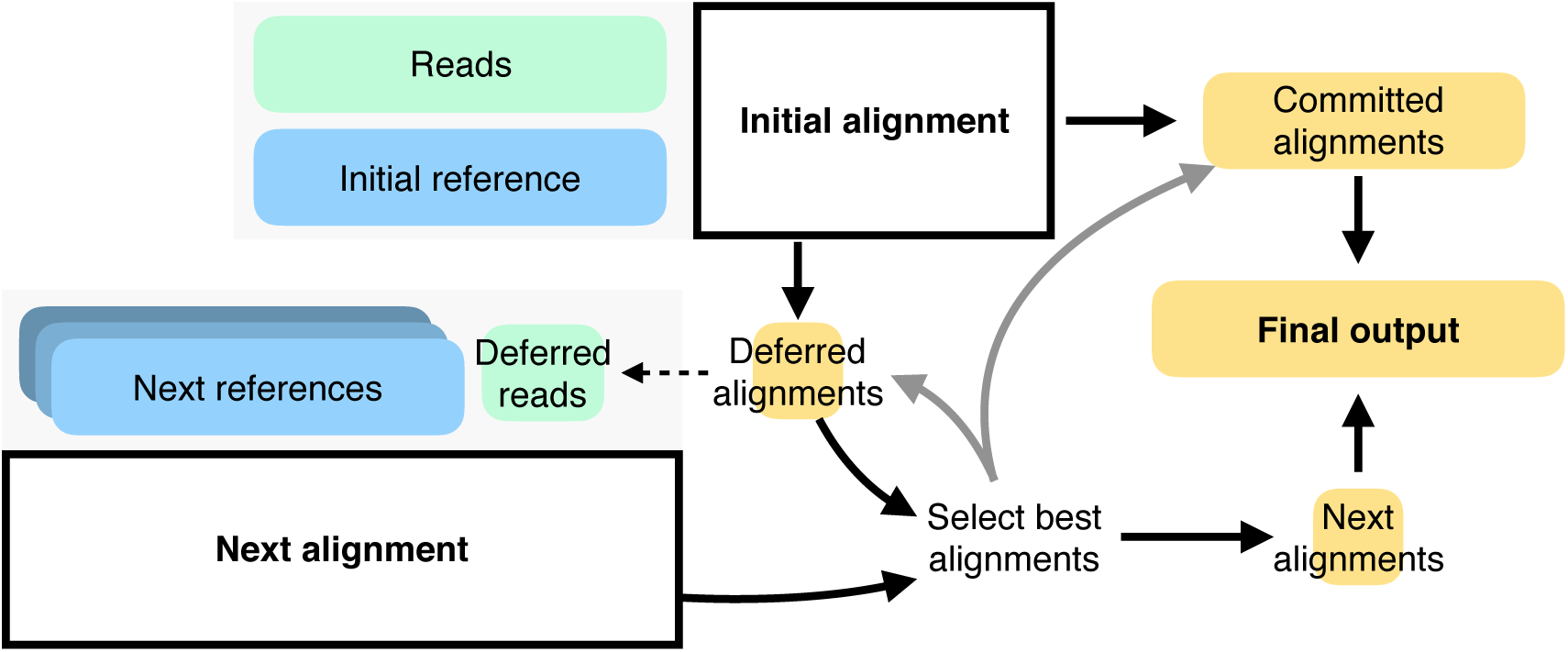
The reference flow workflow: Reads are aligned to reference genome in the first pass. Reads with high mapping quality alignments are “committed.” Unaligned reads or reads with low mapping quality are “deferred” and re-aligned to one or more additional references. The process can iterate, with similar logic for how reads are committed or deferred to another pass. Deferrals could follow the shape of an overall “reference flow graph.” Once all alignments are complete, alignments are merged. For a read aligning to more than one reference, only the best is reported, with ties broken arbitrarily. Alignments are translated (“lifted over”) to the coordinates of a standard reference like GRCh38.

To contextualize the results, a diploid “personalized reference genome” – the genome from which reads are simulated – is used as the ideal reference genome for alignment. By considering the alignments to the diploid personalized reference as a rough upper bound on how well any method can do, we can express results in terms of the degree to which a method closes the gap between the standard linear reference and the personalized reference. When aligning simulated sequence reads, our “RandFlow-LD” method closed 71.82% of the gap in sensitivity on median compared to using GRCh38. Our method also reduced reference bias, reducing by 37% the number of strongly biased sites, and lowering the overall reference to alternate allele (REF-to-ALT) ratio from 1.014 to 1.004. When aligning real whole-genome sequencing reads from NA12878, our method reduced the number of strongly biased heterozygous (HET) sites by 13,332 (34%) and lowered the overall REF-to-ALT ratio from 1.072 to 1.016. It achieves similar gains as the vg graph aligner^11^ in terms of alignment accuracy and reference bias avoidance while using just 14% of the memory and 18% of the CPU time. RandFlow-LD can use a larger set of 26 population-level references (“RandFlow-LD-26”) to achieve lower reference bias than vg, while still running twice as fast.

## 2 Results

### 2.1 Standard and major-allele references

We built a global major-allele reference by modifying the GRCh38 primary reference^1^ to contain the most common allele at each bi-allelic SNV or small indel. Common alleles were determined using the 1000 Genomes Project GRCh38 call set^20^. We call this the “global major” reference. We repeated this process but considering only the five subsets of individuals belonging to the five super populations labelled by the 1000 Genomes Project. We call these “superpop major” references. Table 1 summarizes the variants included in each reference. All references were indexed for use with the Bowtie 2 aligner^21^.

**Table 1:** Number and types of variants included in each major-allele reference of chromosome 21. Superpopulation labels are from the 1000 Genomes Project: AFR (African), AMR (admixed American), EAS (East Asian), EUR (European), and SAS (South Asian).

| Group | # samples | # SNVs | # indels | Longest DEL (bp) | Longest INS (bp) |
| --- | --- | --- | --- | --- | --- |
| Global | 2,548 | 25,437 | 4,006 | 44 | 29 |
| AFR | 671 | 27,542 | 4,195 | 44 | 52 |
| AMR | 348 | 26,424 | 4,175 | 44 | 29 |
| EAS | 515 | 26,394 | 4,114 | 44 | 52 |
| EUR | 522 | 25,884 | 4,132 | 32 | 29 |
| SAS | 492 | 26,748 | 4,200 | 44 | 29 |

### 2.2 Simulations for major-allele reference flow

We studied the efficacy of a strategy we call “MajorFlow,” which starts by aligning all reads to the global major reference. Reads that fail to align or align with low mapping quality are deferred to a second pass where they are realigned to each of the 5 superpop major references. For each read we report the best alignment to any reference. We performed all alignments using Bowtie 2 and default parameters^21^, though the method is not restricted to a particular aligner or set of parameters (Section 3.5).

We performed simulation experiments to compare MajorFlow to baselines that used Bowtie 2 to align to the GRCh38 primary assembly^1^ or to major-allele references. We used Mason2^23^ to simulate reads from GRCh38 chromosome 21 (Section 3.1). Starting from the 1000 Genomes Project GRCh38 call set^20,24^, we randomly selected 100 individuals and built personalized, diploid references for each using phased variant calls (Table S2). We included single nucleotide variants (SNVs) and short insertions and deletions (indels). We simulated 1M reads from both haplotypes (2M total) of each individual. Since allelicbalance measurements require deeper coverage, we also simulated a larger set of 20M reads for 25 of the individuals (Table S2). We also assessed the alignment methods using an ideal, diploid personalized reference genome (Section 3.2). Results using the personalized reference serve as a rough upper bound on what is achievable with references that lack foreknowledge of donor genotypes^17,25,26^. We call this a “rough” upper bound because, while the personalized reference is ideal in that it contains the correct variants, the accuracy of alignment is also affected by tool-specific heuristics. A true upper bound would be hard to obtain, so we settle for the rough upper bound provided by the personalized genome, as in previous work^17,27^.

We measured *sensitivity*, the fraction of input reads that aligned correctly, as well as the fraction that aligned incorrectly. We called an alignment correct if its leftmost aligned base was within ±10 bases of its simulated point of origin (Section 3.3). We also measured *allelic balance* at HET SNVs, where we defined allelic balance as the number of alignments with the REF allele divided by the number with either the REF or ALT allele. We also counted the number of *strongly biased sites*, i.e. those with allelic balance *≤*20% or *≥* 80%. Finally, as an aggregate measure of balance, we measured the *overall REF-to-ALT* ratio totaled over all HET sites (Section 3.4).

The MajorFlow method (“MajorFlow” in Figure 2a) exhibited higher sensitivity than single-reference methods that used the standard reference (“GRC”) or any of the major-allele references (“Major”). If we consider the increase in sensitivity relative to the sensitivity gap between the GRCh38 reference and the ideal personalized reference, MajorFlow’s median sensitivity improvement closed about 51.34% of the gap. In terms of number of incorrect alignments, MajorFlow closed 46.81% of the benefit of personalization (Figure S15). MajorFlow’s sensitivity was still higher when we enhanced the major-allele strategy by always matching the ethnicity of the major-allele reference to that of the simulated sample (“Matched”). Alignments for reads simulated from the African super population (AFR) had lower sensitivity compared to the others, even when aligned to the AFR superpop-major reference (Figure S1). This is consistent with AFR’s greater genetic heterogeneity due to the out-of-Africa bottleneck. Consistent with past results, there was only a small difference in mapping sensitivity when using the global-major versus the superpop-major references, even when the simulated donor’s ethnicity was matched with the reference (Figure 2a).

**Figure 2:**
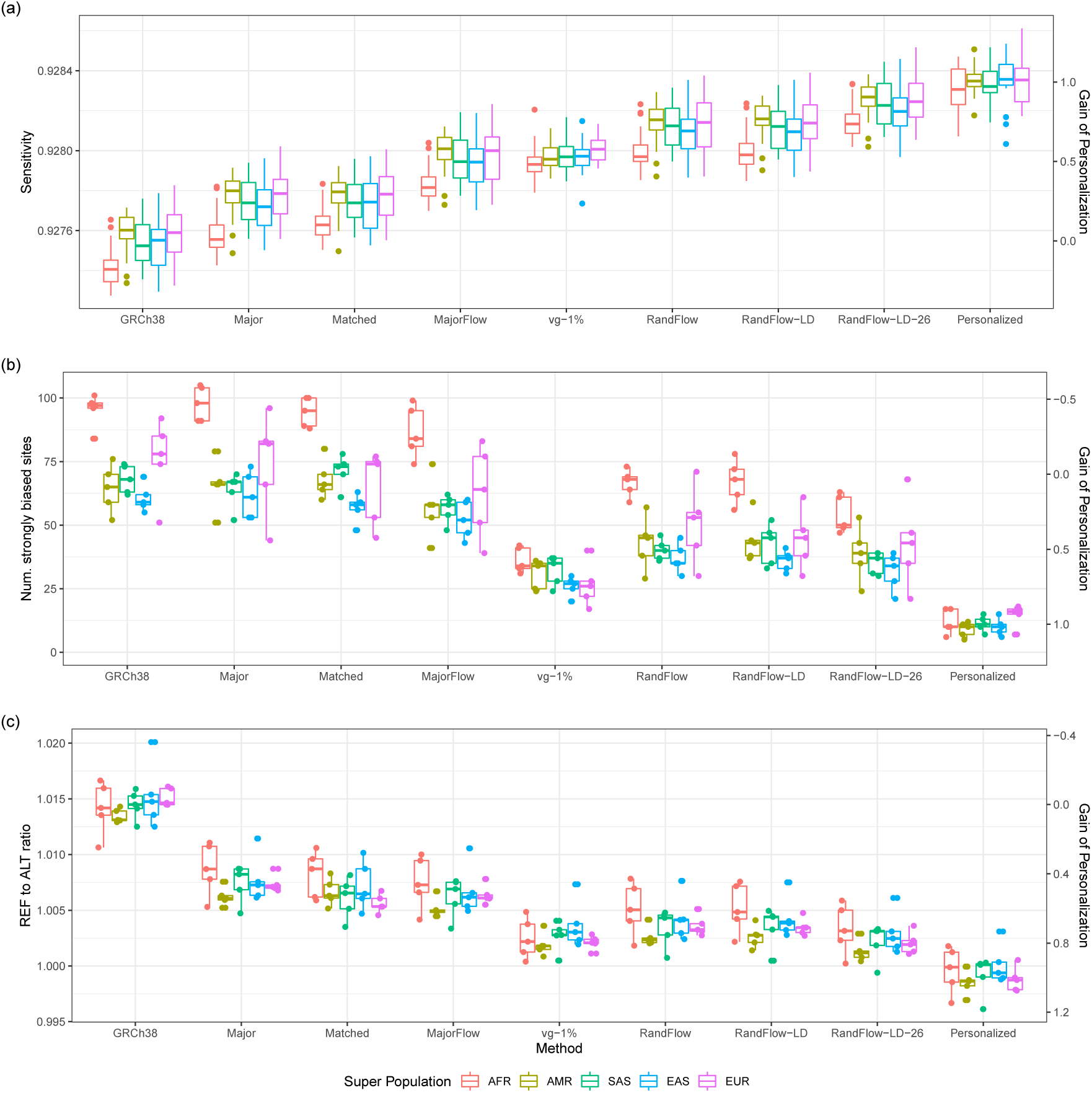
Alignment results using different methods. (a) Alignment sensitivity for 100 samples selected from the 1000 Genomes Project; 2 million reads are simulated from each sample. (b) The number of strongly biased heterozygous sites, and (c) the overall REF-to-ALT ratio for 25 samples; 20 million reads are simulated for each sample. The columns are sorted by median alignment sensitivity.

MajorFlow also reduced reference bias relative to the single linear references using the set of 25 deeper simulations. Overall REF-to-ALT ratio decreased from 1.0145 using the standard reference to 1.0073 using the global major reference, then further to 1.0064 using MajorFlow method (Figure 2c). The median number of strongly biased HET sites dropped from 70 for GRCh38 to 59 for MajorFlow (Figure 2c).

### 2.3 Simulations for stochastic reference flow

While MajorFlow outperformed the single-linear-reference strategies, we noticed it was less effective than the graph-based vg aligner at increasing sensitivity or reducing reference bias (Figure 2). We hypothesized this was because the major-allele references used by MajorFlow were too similar to each other, narrowing the genetic diversity visible to the method. Specifically, the mean edit distance between all pairs of superpop major references was 15,115 bp for chromosome 21, whereas the mean between all pairs of five individuals randomly drawn from the super populations was 47,966 bp (Figure S4).

We designed two alternative methods that draw on super-population-specific variation while also keeping the second-pass genomes genetically distinct. “RandFlow” generates a random reference haplotype for each super population by performing an independent draw at each variant site, choosing the ALT allele with probability equal to its frequency in the super population. “RandFlow-LD” is similar but additionally maintains some linkage disequilibrium (LD). RandFlow-LD begins by choosing one haplotype from the super population uniformly at random. Then, starting at the first (leftmost) polymorphic site in the haplotype and for the length of a 1,000-bp window, it selects alleles matching the chosen haplotype. At the next polymorphic site beyond the 1,000-bp window, the method chooses a new super-population haplotype uniformly at random and repeats the process. In this way, variant selection is still weighted by allele frequency (since haplotypes are selected uniformly at random) but a degree of LD is also maintained. Both strategies result in greater genetic distances between the second-pass references compared to MajorFlow, with mean pairwise distances on chromosome 21 being 47,316 for the Rand-Flow strategy and 46,326 for RandFlow-LD. Further details are in Section 3.5.

Using the chromosome-21 simulation data from the previous section, we observed that RandFlow and RandFlow-LD achieved higher sensitivity and lower numbers of incorrect alignments compared to MajorFlow. If we consider the increase relative to the sensitivity gap between the GRCh38 reference and the ideal personalized reference, RandFlow’s and RandFlow-LD’s median sensitivity improvement closed about 70.91% and 71.82% of the gap respectively (Figure 2a). The reduction of incorrect alignment compared to personalization was 66.22% for RandFlow and 67.34% for RandFlow-LD (Figure S15). While RandFlow slightly underperformed RandFlow-LD in sensitivity and number of incorrect alignments, we note that RandFlow does not require that variants be phased, and so can benefit from larger compendia of unphased genotypes available through projects like gnomAD^28^.

Using the set of 25 deeper simulations, RandFlow-LD reduced the median number of strongly biased HET sites to 44, from a median of 70 using the GRCh38 reference. RandFlow-LD also reduced the overall REF-to-ALT ratio to 1.0038, an improvement over GRCh38 (1.0145) and MajorFlow (1.0064).

We further compared the reference flow methods to vg^11^. vg aligns to a reference that is shaped as a graph rather than a string. Paths through the graph correspond to different combinations of REF and ALT alleles. Such methods can improve alignment accuracy and reduce reference bias by adding paths — thereby removing alignment-score penalties — for ALT alleles. We built a vg index using chromosome 21 of the GRCh38 primary assembly as the base, and including all variants from the 1000-Genomes GRCh38 callset having allele frequency at least 1% and aligned all reads to the graph. There were 192,846 variants passing the threshold, about twice as many ALT alleles as we considered in our RandFlow (93,146) and RandFlow-LD (95,319) strategies. We found that RandFlow and RandFlow-LD had higher sensitivity and fewer incorrectly aligned reads than vg (Figures 2a and S15), but that vg yielded a smaller number of strongly biased sites (30, versus 44 for RandFlow-LD) and a slightly more balanced overall REF-to-ALT ratio (1.0026, versus 1.0038 for RandFlow-LD). While neither approach is the clear winner in this comparison, the reference flow methods use substantially less time and memory, as discussed below.

To explore how using more second-pass genomes improves accuracy, we used the same RandFlow-LD method to make a set of 26 population-specific chromosome 21 sequences. These correspond to the 26 separate populations studied in the 1000 Genomes Project, subdividing the 5 super populations and including 168,593 variants in total. The alignment sensitivity of this “RandFlow-LD-26” approach was the best of any we evaluated, closing 84.08% of the gap between the GRCh38 and personalized references. It achieved lower allelic bias compared to RandFlow-LD, with a median of 39 strongly biased sites and an overall REF-to-ALT ratio of 1.0024. Though it used a total of 27 references (including the first-pass major-allele reference), RandFlow-LD-26 used less CPU time and had a smaller memory footprint compared to vg (Section 2.5).

### 2.4 Assessing reference bias with real data

We further assessed these methods using a deep whole-genome sequencing dataset from individual NA12878 (SRR622457) generated by the 1000 Genomes Project^20^. The dataset consisted of 1.4 billion Illumina HiSeq 2000 101-bp paired-end reads, though we used only the first end of the pair in these experiments. Since each read’s true point of origin is unknown, we assess only allelic balance and not sensitivity. We assessed allelic balance only at sites where NA12878 is HET according to the 1000 Genomes Project GRCh38 call set, and then stratified the sites according to the Genome-in-a-Bottle v3.3.2 confidence annotation^29^. There were 1,723,317 (83%) HET sites in high-confidence regions, and 344,945 (17%) in low-confidence regions. We also constructed and aligned to an ideal, diploid personalized reference using the phased variant calls for NA12878 from the GRCh38 call set. We assessed only the RandFlow-LD and RandFlow-LD-26 methods since they outperformed other reference-flow methods in the simulation experiments. After a first-pass alignment to the global major-allele reference, there were 250M (17.4%) reads deferred into the second pass.

Consistent with the simulation experiments, we observed that RandFlow-LD and vg both reduced the number of strongly biased sites in all regions, from 44,810 in the case of GRCh38, to 34,429 (23% reduction) for RandFlow-LD and 31,784 (29% reduction) for vg (Figure 3 and Table 2). Similarly, RandFlow-LD reduced the overall REF-to-ALT ratio from 1.0719 (GRCh38) to 1.0160 and vg reduced it to 1.0123. Further, RandFlow-LD-26 reduced the number of strongly biased sites to 30,317 (32% reduction) and REF-to-ALT ratio to 1.0081, best among the methods using non-personalized references. The variant-aware methods substantially reduced reference bias compared to a method that aligned only to the global major reference (“Major”). In high-confidence regions, variant-aware methods reduced the number of strongly biased sites by 39% – 50% compared to GRCh38, and reduced the REF-to-ALT ratios from 1.041 to about 1.01 (Figure S17). In low-confidence regions, we observed 11% – 18% reduction in number of strongly biased sites, but a greater benefit in REF-to-ALT ratios, from 1.024 to 1.001–1.028 (Figure S18). RandFlow-LD-26 reduced bias most among variant-aware approaches.

**Table 2:** Measures of allelic balance for NA12878 whole genome sequencing dataset stratified by Genome-in-a-Bottle v3.3.2 confidence annotation. vg index includes variants with 10% or higher allele frequency in the 1000-Genomes Project GRCh38 call set. This methods are sorted by REF-to-ALT ratio in all regions.

| Method | REF-to-ALT ratio | Total # biased | # biased toward REF | # biased toward ALT |
| --- | --- | --- | --- | --- |
| **High confidence** |  |  |  |  |
| GRCh38 | 1.0407 | 20,012 | 18,141 | 1,871 |
| Major | 1.0227 | 19,837 | 15,415 | 4,422 |
| RandFlow-LD | 1.0133 | 12,239 | 9,512 | 2,727 |
| vg | 1.0124 | 10,518 | 7,971 | 2,547 |
| RandFlow-LD-26 | 1.0098 | 9,984 | 7,489 | 2,495 |
| Personalized | 1.0033 | 7,024 | 4,600 | 2,424 |
| **Low confidence** |  |  |  |  |
| GRCh38 | 1.2355 | 24,798 | 20,594 | 4,204 |
| Major | 1.1230 | 26,579 | 20,108 | 6,471 |
| RandFlow-LD | 1.0282 | 22,190 | 15,891 | 6,299 |
| vg | 1.0120 | 21,266 | 15,422 | 5,844 |
| RandFlow-LD-26 | 1.0008 | 20,333 | 13,874 | 6,459 |
| Personalized | 0.9750 | 16,266 | 9,299 | 6,967 |
| **All regions** |  |  |  |  |
| GRCh38 | 1.0718 | 44,810 | 38,735 | 6,075 |
| Major | 1.0397 | 46,416 | 35,523 | 10,893 |
| RandFlow-LD | 1.0160 | 34,429 | 25,403 | 9,026 |
| vg | 1.0123 | 31,784 | 23,393 | 8,391 |
| RandFlow-LD-26 | 1.0081 | 30,317 | 21,363 | 8,954 |
| Personalized | 0.9981 | 23,290 | 13,899 | 9,391 |

**Figure 3:**
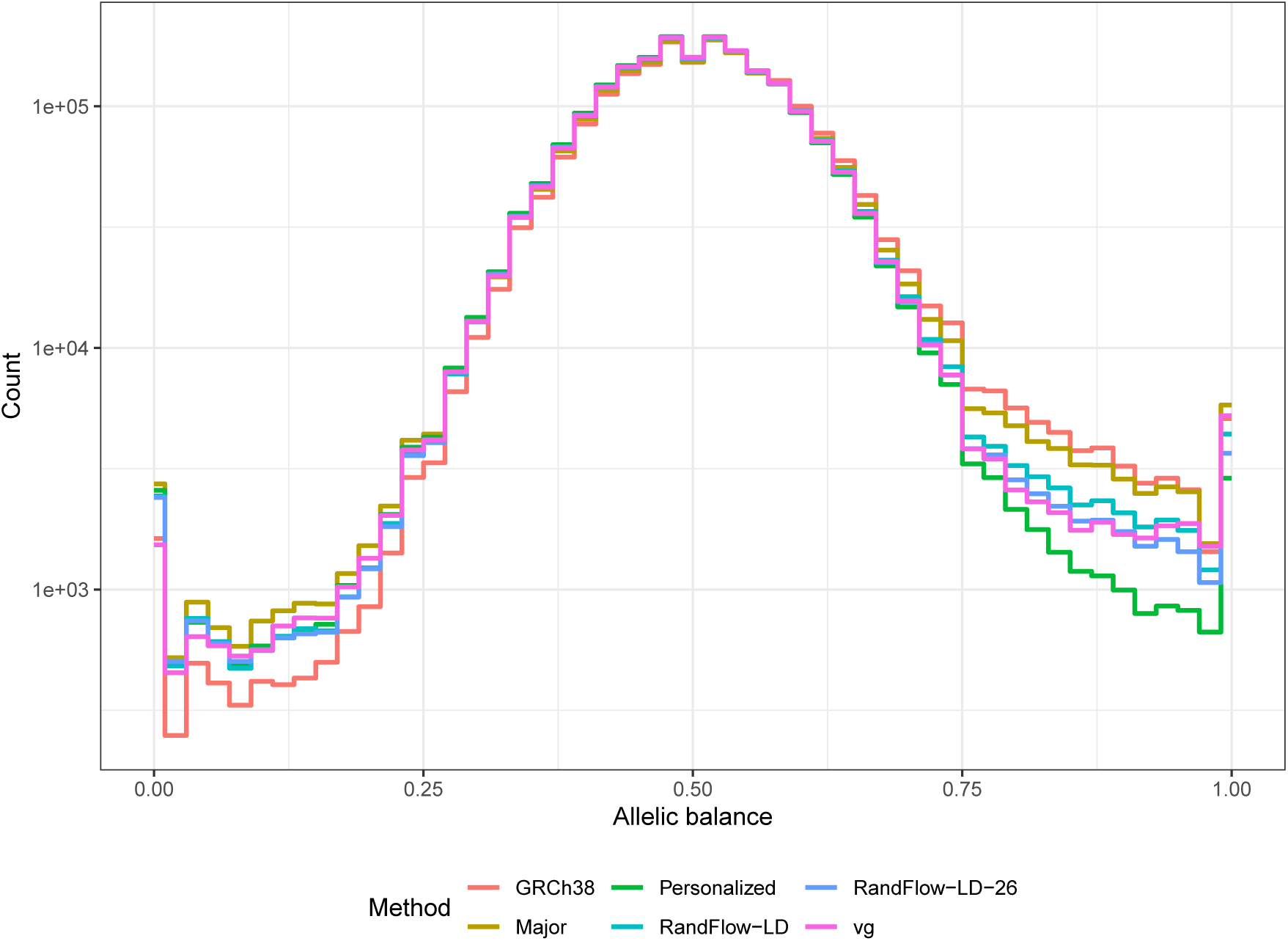
Histograms of allelic balance using a high-coverage real WGS dataset of individual NA12878 (SRR622457). Experiments are performed using GRCh38 (*GRC*), global major reference (*Major*), diploid personalized genome (*Personalized*), vg using alleles with frequency ≥ 10% (*vg*), reference flow using 1000-bp phased blocks with 5 super populations (*RandFlow-LD*) and reference flow using 1000-bp phased blocks with 26 populations (*RandFlow-LD-26*).

Notably, the number of strongly biased sites was still as high as 23,290 when aligning to an ideal personalized reference (Table 2). In part this is because the 1000 Genomes Project calls include only a subset of the variation present in the actual NA12878 genome. This is both because some genomic regions were excluded from the call set because of low mappability, and because the call set does not include larger-scale structural variants that can have an outside effect on sensitivity and bias. We also noted that the more strongly biased sites were biased toward REF (13,899) more often than toward ALT (9,391) when aligning to the personalized reference, supporting the argument that variants missing from the call set are affecting the bias.

To better understand where variant-aware methods reduce bias the most, we studied the relationship between highly biased HET sites and various categories of repeat families (Figure 4) and classes (Figure S5) annotated by RepeatMasker^30^. Using alignment to GRCh38, many strongly biased HETs are in L1 (10,288, or 23%) and Alu (11,255, or 25%). RandFlow-LD greatly reduced the number of strongly biased HET sites in L1 (to 5,250, reduced by 49%) and Alu (to 6,555, reduced by 42%). A similar reduction is observed when using vg, but the greatest reductions are achieved by RandFlow-LD-26. For instance, RandFlow-LD-26 reduces the number of strongly biased sites in L1 from 10,288 to 3,560, a 65% reduction.

**Figure 4:**
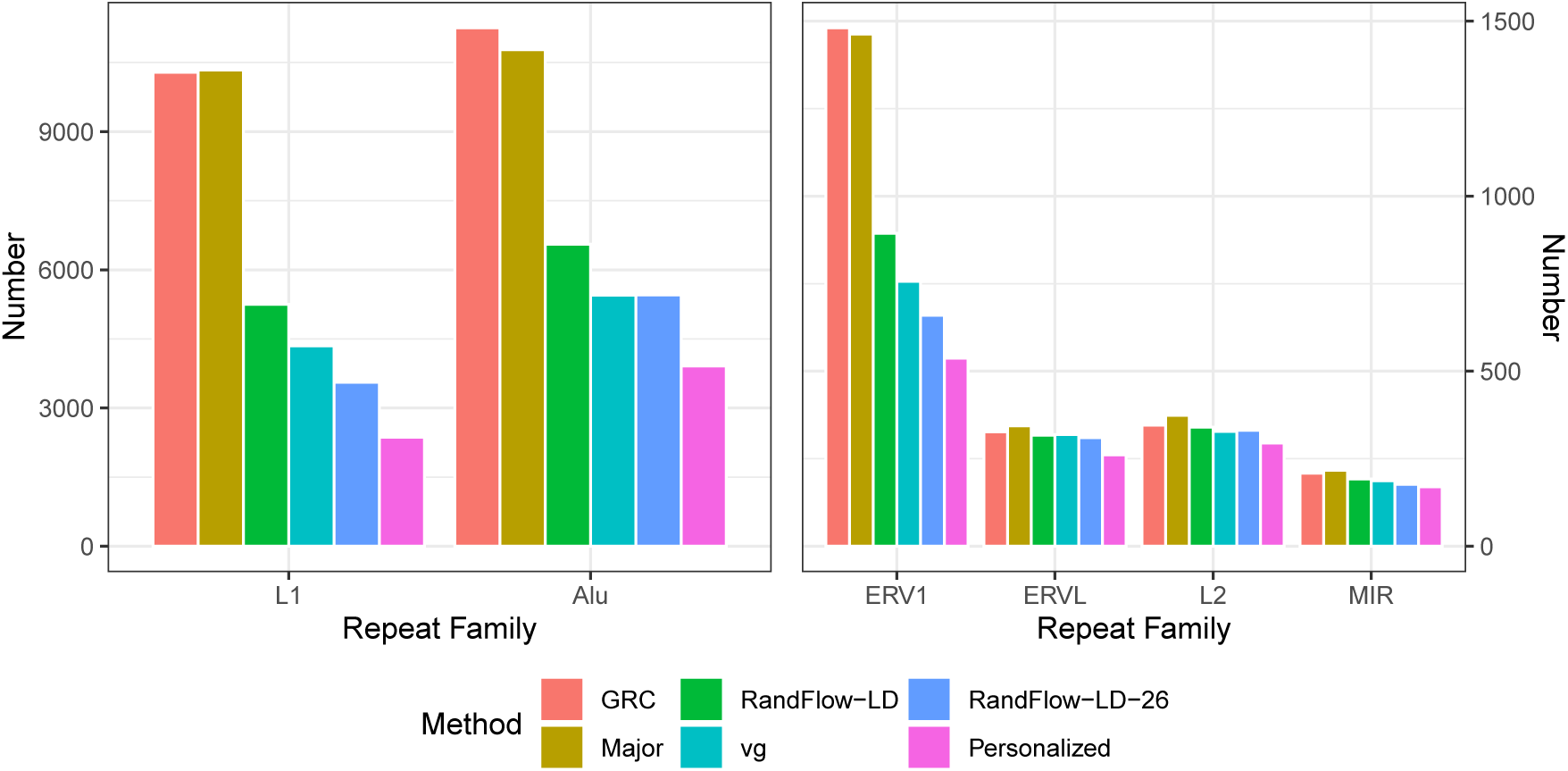
Number of strongly biased HET sites stratified by RepeatMasker class, after aligning single-end reads from SRR622457. HET sites are determined using 1000 Genomes Project calls for NA12878, the individual sequenced in SRR622457. RandFlow methods and vg reduce the number of biased sites substantially for L1, Alu and ERV1. RandFlow-LD-26 reduces the number of biased sites most among the methods tested.

### 2.5 Computational efficiency

We constructed a dataset consisting of 10M single-end reads randomly sampled from the first end of the SRR622457 paired-end dataset. We ran each alignment method and measured the total size of index files on disk, the peak memory usage, and the CPU time (Table 3). We measured peak memory usage using the maximum resident set size reported by the GNU Time utility. We also measured CPU time using GNU Time. We performed the experiments on a computer with a 2.2 Ghz Intel Xeon CPU (E5-2650 v4) and 515GB memory. We configured all read-alignment jobs to use 16 simultaneous threads but otherwise left parameters at their defaults. Though RandFlow-LD and RandFlow-LD-26 were the only reference-flow approaches we benchmarked here, we expect MajorFlow and Rand-Flow to perform similarly to RandFlow-LD since they execute the same sequence of steps, using the same number of linear reference genomes.

**Table 3:**
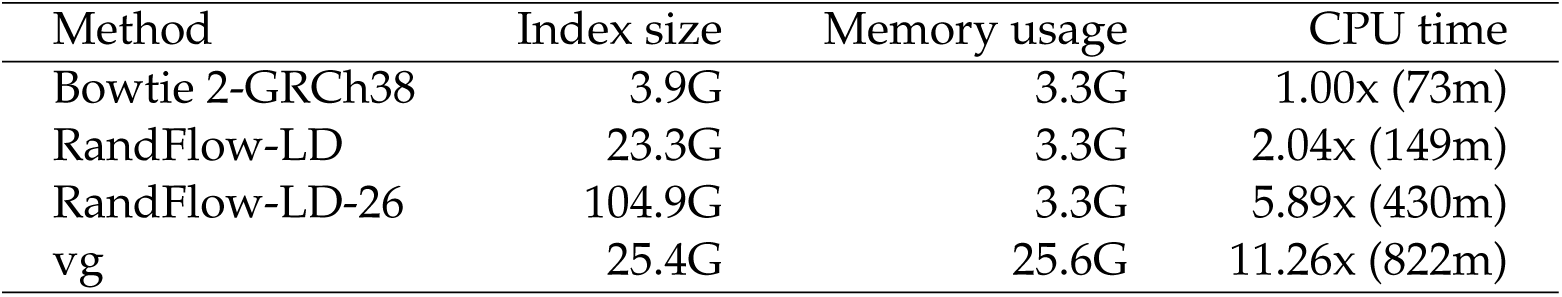
Comparison of alignment methods using 10M single-end 101-bp reads from individual NA12878 (SRR622457). The vg index includes variants with allele frequency ≥ 10% in the 1000-Genomes Project GRCh38 call set. The RandFlow-LD indexes include the indexes for liftover and Bowtie 2 indexes for the global major-allele reference as well as second-pass population references. CPU time is compared to a baseline run of Bowtie 2 to the GRCh38 primary assembly.

| Method | Index size | Memory usage | CPU time |
| --- | --- | --- | --- |
| Bowtie 2-GRCh38 | 3.9G | 3.3G | 1.00x (73m) |
| RandFlow-LD | 23.3G | 3.3G | 2.04x (149m) |
| RandFlow-LD-26 | 104.9G | 3.3G | 5.89x (430m) |
| vg | 25.4G | 25.6G | 11.26x (822m) |

Compared to an alignment run against the GRCh38 primary assembly, RandFlow-LD used about 5.97 times as much disk space to store the reference index files, consistent with the fact that RandFlow-LD uses 1 reference in the first pass and 5 in the second (Table 3). vg used a similar amount similar size of disk space for its indexes (.xg, .gcsa and .gcsa.lcp). vg used 7.31 times as much peak memory usage compared to the linear-based alignment methods, including RandFlow-LD. The baseline approach used less than 9% of the CPU time as vg, while RandFlow-LD used less than 18% of the CPU time as vg. Overall, RandFlow-LD used only about twice as much CPU time as the baseline. 84% of RandFlow-LD’s runtime overhead was spent in re-alignment, 13% was spent in liftover and less than 2% was spent in merging alignments. When extending RandFlow-LD to RandFlow-LD-26, the CPU time increased to 589% of the baseline and the index size increased to 104.9 GB. But its speed was 1.9 times that of vg.

We note that RandFlow-LD and RandFlow-LD-26 have similar peak memory footprint to the baseline because the reference-flow software runs the alignment jobs serially. In other words, only one reference genome index is resident in memory at a time. Because the read aligners themselves are multithreaded, we can do this while using many simultaneous threads.

### 2.6 Comparison of variant-aware alignment approaches

We further compared the reference flow methods with other graph-based methods, including the graph aligner HISAT2^12^ (Figures S6, S7 and S8). HISAT2 was computationally efficient, using 46.5% of the CPU time compared to Bowtie2 with a whole-genome graph containing variants with allele frequency *≥* 10% in the 1000 Genomes Project. Its index size (6.1G) and memory usage (6.5G) were small compared to vg’s (index size: 25.6G; memory usage: 25.6G) using the same variant set. However, it performed worse than other methods on mapping sensitivity (92.46%, versus 92.80% for vg), median number of strongly biased sites (138, versus 30 for vg) and overall REF-to-ALT ratio (1.0265, versus 1.0026 for vg) when evaluated using simulated reads from chromosome 21.

To understand the effect of including different numbers of variants in the vg graph, we tested a few vg graph sizes: a vg graph with no variants (just the linear GRCh38 reference), a graph with all 1000-Genomes variants having *≥*10% allele frequency (AF), and a graph with all *≥*1% AF variants. For a more direct comparison with RandFlow-LD, we also made a vg graph that included the union of the variants used in all RandFlow-LD references (Note S1, Figure S5). We indexed the graphs and evaluated alignment performance using the same simulation framework as in sections 2.2 & 2.3. The median mapping sensitivity of the 10% AF graph outperforms other vg-based methods (*≥*10% AF: 92.805%; *≥*1% AF: 92.797%), while the ≥1% AF graph gave fewer median strongly biased sites (≥ 10% AF: 40; ≥ 1% AF: 30) and lower overall REF-to-ALT ratio (*≥*10% AF: 1.0051; ≥ 1% AF: *≥*1.0026). When comparing RandFlow-LD with the vg graph built using the RandFlow-LD variants (*vg-RandFlow-LD* column in Figures S6, S7 and S8), RandFlow-LD is more sensitive (92.82%, versus 92.80% for *vg-RandFlow-LD*), achieves a more balanced REF-to-ALT ratio (1.0038, versus 1.0069 for *vg-RandFlow-LD*), and yields a smaller number of highly biased sides (44, versus 50 for *vg-RandFlow-LD*).

## 3 Methods

### 3.1 DNA data simulation

We built diploid consensus genomes for the selected individuals (Table S2) using bcftools31 based on the SNVs and indels specified by the 1000 Genome Project GRCh38 call set^24^. We used Mason2^23^ to simulate paired-end Illumina 100-bp reads, but used only the first end in most experiments. Since variants were already included in the reference genomes we simulated from, we did not use Mason2’s variation-adding feature. We enabled Mason2’s features for generating random sequencing errors and quality values. We simulated reads independently from each haplotype to generate diploid read sets, keeping information about the haplotype, chromosome and offset of origin for downstream evaluations.

### 3.2 Building and aligning to the personalized reference

We built personalized, diploid reference genomes for each of the 100 randomly selected 1000 Genomes individuals^5,32^ (Table S2). We used phased variant calls – including SNVs and indels and including sites with more than 2 ALT alleles – from both haplotypes of the selected individual to build FASTA files containing a personalized diploid reference genome. When aligning to the personalized diploid references, we aligned all reads separately to both haplotypes. We aligned to the haplotypes separately so that the mapping qualities could be informative; aligning to both together would have yielded consistently low mapping qualities. We then merged the resulting alignments. For a read that aligned to both haplotypes, we took the alignment with the higher alignment score. We broke ties by taking the alignment with higher mapping quality or, if the tie remained, at random.

For the simulated experiment using chr21, we aligned to each personalized haplotype 5 separate times, providing the aligner with 5 different random seeds. This yielded 10 total alignments from which we selected the best. This helped to improve the upper bound somewhat, since the 5 random seeds gave the aligner 5 times as many chances of finding the best alignment even with the censoring effect of alignment heuristics (Figure S6).

### 3.3 Measuring sensitivity

In simulation experiments, we keep information about each read’s haplotype, chromosome and offset of origin. We say a read aligns correctly if the alignment’s leftmost mapped base is within *±*10-bp of the leftmost base at the read’s point of origin. Since we use Bowtie 2 with default alignment parameters, no “soft clipping” is possible and it does not affect the definition of correctness. Reads that align outside of the *±*10-bp window are called incorrect. We define *sensitivity* as the fraction of reads that aligned correctly.

### 3.4 Allelic balance measurement

We measured allelic balance at each bi-allelic HET SNVs reported in the 1000 Genomes Project GRCh38 call set. HET SNVs that were contained within a larger deletion variant were excluded, whether or not the deletion was heterozygous. At each relevant HET, we considered the “pileup” of alleles at the site induced by overlapping read alignments. Let *a*ref and *a*alt denote the number of REF and ALT alleles overlapping the site:

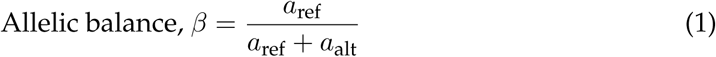

We say a site is *strongly biased* when *β ≤* 0.2 or *β ≥* 0.8. For a collection of sites, we calculate the *overall REF-to-ALT ratio* as total number of REF alleles divided by the total number of ALT alleles across the sites:

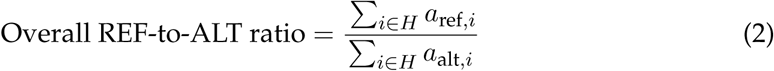

We ignore alleles besides REF and ALT, and we ignore alignments having a gap at the site. The assumption that on average *β* should equal 0.5 at HET sites is well founded for simulated datasets. Real datasets have biases, which might be due to systematic sequencing errors or fragmentation bias, for example. Biases might also arise from errors in the set of sites we consider to be HET, e.g. if the variant caller that produced the HET calls was itself affected by allelic bias.

### 3.5 Reference flow

#### Preparation

The reference-flow methods require that we first build read-alignment indexes and coordinate-translation indexes for the relevant species and populations. Both can be generated from a reference genome in FASTA format and a collection of population variants in VCF format. The reference-flow software (a) processes the VCF to select variants to include in the population reference genomes, (b) generates both the first-pass and the second-pass references based on the reference genome, and (c) builds Bowtie 2 indexes for all references.

For convenience, we provide pre-built RandFlow-LD genomes and indexes based on the GRCh38 reference and the 1000 Genomes Project GRCh38 call set (see Availability of data and materials).

#### First pass

In the first pass, we align all reads to an initial reference genome. For the particular reference-flow strategies evaluated here (MajorFlow, RandFlow, RandFlow-LD, and RandFlow-LD-26), we first aligned to the “global major” reference (Section 2.2). Reads that fail to align or that align with low mapping quality are “forwarded” to a second pass, whereas reads that align with high mapping quality are “committed” and are ultimately passed through to the final output. We use a mapping-quality threshold because it is readily available – reported by most popular read aligners – and because alignments with low MAPQ are the most likely to benefit from the second alignment pass. After empirical experiments, we selected a MAPQ threshold of 10 (Figures S2 and S13).

#### Second pass

For reads forwarded to the second pass, we realign to a set of references that include a wider range of genetic variation. In the methods evaluated here other than RandFlow-LD-26, we use five second-pass references, each corresponding to a 1000 Genomes Project superpopulation: AFR (African), AMR (admixed American), EAS (East Asian), EUR (European), and SAS (South Asian). For RandFlow-LD-26, we use 26 second-pass references, each corresponding to a population in the 1000 Genomes Project. In the case of the MajorFlow method, the second-pass genomes are simply the major-allele references corresponding to each of these superpopulations (Section 2.2). In all cases, the second-pass references consist of a single haplotype.

#### Stochastic references

In the RandFlow, RandFlow-LD and RandFlow-LD-26 strategies, second-pass references are designed to represent “random individuals” from the super populations. For RandFlow, we construct the second-pass references by iterating through each polymorphic site *i* and performing an independent random draw to choose the ALT allele with probability equal to its allele frequency *pi* in the super population:

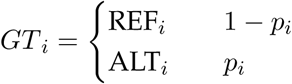

In the case of the RandFlow-LD and RandFlow-LD-26 strategies, for a variant site we select one haplotype in the super population uniformly at random. We then maintain the linkage disequilibrium (LD) relationship by selecting the genotypes from the same haplotype for the next 1000-bp region.

While we used the population and super population labels provided by the 1000 Genomes Project here, the reference-flow framework can work with any granularity of label. Further, neither the MajorFlow nor the RandFlow strategies require that genetic variants be phased. Those approaches could also work with larger, unphased compendia of genetic information such as GnomAD^28^.

#### Merging and lifting

For reads that aligned to more than one reference, we must choose a single “best” alignment to include in the ultimate SAM output. We select by choosing the alignment with the highest alignment score; roughly, this corresponds to the alignment with the fewest mismatches and gaps. If there is a tie for best alignment score, the alignment with higher mapping quality is selected. If there is a tie in both categories, we select at random from among the tied alignments.

For maximum compatibility with downstream tools, the SAM output from our reference-flow methods is with respect to the standard GRCh38 primary assembly. But since the reference genomes in our method – including the major-allele references – can have insertions or deletions with respect to the standard reference, we must translate (“lift over”) these alignments to standard reference coordinates before outputting them. We implemented a simple method to lift over alignments that builds a succinct mapping of coordinates from a genome to the standard reference genome using a VCF file. We use the mapping to adjust the POS and CIGAR fields of a SAM file so as to be compatible with the standard reference. The time and memory used to lift the alignments were included in the benchmarking measurements discussed in Section 2.5.

## 4 Discussion

We proposed and evaluated a family of “reference-flow” alignment methods. These are based on the idea that reads that fail to align or align poorly to one reference might align well to another with a different complement of alleles. We first showed that a 2-pass method using superpopulation major-allele references (MajorFlow) outperformed both a standard linear reference and individual major-allele references. As a further improvement, we proposed the RandFlow and RandFlow-LD methods that align to “random individuals” from each super population. These methods performed similarly to vg and approached the performance achieved using the ideal, personalized reference. The reference flow methods were much more computationally efficient than vg, running 5.5 times as fast and using 14% of the memory compared to vg when aligning to a graph containing all 1000 Genomes variants of frequency 10% or higher.

Our results complement key points from previous studies. Like the FORGe study^17^, we also showed that alignment to a major-allele reference improves alignment accuracy compared to the standard linear reference. Also like FORGe, we showed that aligning to a super-population-matched major-allele reference did not substantially improve alignment accuracy compared to a global major-allele reference combining all super populations. Our results also reinforce that a linear aligner can be extended to incorporate variants and exhibit similar accuracy to a graph aligner^16,33^.

For compatibility with downstream tools, alignments output by reference-flow methods must have their reference coordinates translated back to the standard linear reference. Notably, this requires only a pairwise alignment from each of the reference-flow references to the standard reference. Thus, approaches such as RandFlow and RandFlow-LD use 5 references in the second pass require 6 pairwise whole-genome alignments: one from the first-pass major-allele reference to the standard reference, and 5 from each of the second-pass references. This can be advantageous in the situation where the reference-flow genomes are assemblies with no pre-existing multiple alignment (e.g. VCF file) describing their relationship. Algorithms for calculating genome-scale multiple alignments are resource intensive^34,35^ and yield a more complex structure compared to a pairwise alignment. Reference flow’s use of pairwise alignments also helps to solve an “N+1” problem; adding one additional reference to the second pass requires only that we index the new genome and obtain an additional whole-genome alignment (or otherwise infer such an alignment, e.g. from a VCF file) to the standard reference. We demonstrated that we could extend reference flow to 26 1000-Genomes populations, reducing bias still further while still aligning faster than vg. This flexibility could be important in the coming era where long and ultra-long sequencing reads allow us to build many high quality human genome assemblies.

While we explored methods involving a single initial reference and a set of second-pass references based on 1000-Genomes populations or super populations, we can also consider a wider class of possible architectures. For instance, considering that our method consistently performs worst on the AFR super population, we could imagine building a deeper “tree” of AFR-covering references. A read aligning poorly to the second-pass reference representing the AFR super population could, in a third pass, be aligned to an array of references for specific populations within AFR. We can imagine more complex architectures as well, leading to a general notion of a “reference flow graph” where nodes represent references and directed edges indicate which references might be attempted next. Whether a read should be forwarded along an edge would be dictated by a (possibly edge-specific) rule that uses alignment score, mapping quality, whether the alignment overlapped some genomic region, or other factors.

Our approach for selecting population-specific genomes involves randomness, chiefly as a way of “pushing” genomes further apart compared to the major-allele references. An alternative would be to cast this as a problem of optimizing the references’ “coverage” of the overall genotype space. Such an optimization approach might improve coverage (and therefore accuracy) while removing the random element. This might be accomplished using unsupervised, sequence-driven clustering methods^36,37^, using the “founder sequence” framework^38,39^, or using some form of submodular optimization^40^. A more radical idea is to simply index all available individuals, forgoing the need to choose representatives; this is becoming more practical with the advent of new approaches for haplotype-aware path indexing^33^ and efficient indexing for repetitive texts^41^.

Since reference flow is essentially a “wrapper” that can be placed around an existing aligner, Bowtie 2 could be replaced by a different linear aligner such as BWA-MEM or even a graph aligner such as vg. It is even possible for different nodes in the graph to use different alignment tools. Since the wrapper is written using Snakemake^42^, it is easily deployable both in one-sample single-computer scenarios and in scenarios involving many samples or a collection of networked computers.

In the future, it will be important to benchmark reference-flow methods when larger structural variants are included in the references. Structural variants have a disproportionate effect on alignment quality^43,44^. In principle they are not difficult to include in the reference flow framework, though our lift over procedure is not currently robust enough to handle more complex structural variants like inversions or rearranges.

## 5. Acknowledgments

Part of this research project was conducted using computational resources at the Maryland Advanced Research Computing Center (MARCC). Reference-flow indexes are made freely available on Amazon Web Services thanks to the AWS Public Dataset Program.

## 7. Availability of data and materials

The reference flow software is available at: https://github.com/langmead-lab/reference_flow under the open source MIT license. The experiments described in this paper are available at: https://github.com/langmead-lab/reference_flow-experiments under the open source MIT license. Pre-built RandFlow-LD genomes and indexes based on the GRCh38 reference and the 1000 Genomes Project GRCh38 call set are available at: https://genome-idx.s3.amazonaws.com/bt/flow/randflow_ld.tar.gz. A similar package tht uses the RandFlow-LD-26 references instead of the RandFlow-LD references is available at: https://genome-idx.s3.amazonaws.com/bt/flow/randflow_ld_26.tar.gz. Software versions used in the experiments are specified in Table S1.

## 8. Authors’ contributions

NC, BS, and BL designed the method. NC and BS wrote the software and performed the experiment. TM wrote the liftover software. SI designed and performed the reference bias experiment. NC, BS, TM, and BL wrote the manuscript. All authors read and approved the final manuscript.

## 9. Ethics approval

Not applicable.

## 10. Consent for publication

Not applicable.

## 12. Additional Files

### 12.1 . Additional file 1 — Supplementary information

Contains Note S1–S2, Tables S1–S5, and Figures S1–S18. PDF, 425 KB.

