## Supplementary Information for "Reducing reference bias using multiple population reference genomes"

### Supplementary Notes

#### S1 RandFlow-LD variant set

We built a vg graph using the variant set selected by RandFlow-LD, including the major alleles and five superpop genomes, using `bcftools merge`. This alternative vg graph contains 6,471,383 variants across the whole GRCh38 (Table S5). The size of this set is comparable to the number of variants with $\geq$ 10% allele frequency and is less than one half of the number of variants with $\geq$ 1% allele frequency in the 1000 Genomes Project.

We further compared the allele frequency distribution of chromosome 21 variants in the RandFlow-LD set and the allele frequency $\geq$ 10% set (Figure S9). Because the RandFlow-LD method includes a major-allele reference, all variants with allele frequency greater than 50% are included in the set. There are 14,699 (15%) RandFlow-LD variants with allele frequencies in range $[0.1, 0.2)$, lower than 25,294 (28%) of the allele frequency $\geq$ 10% set. One fifth (19,279) of the RandFlow-LD variants have allele frequencies lower than 10%.

#### S2 RandFlow-LD variability

We assessed the variability of the RandFlow-LD method when using different random seeds. We generated 15 set of RandFlow-LD chr21 references, each containing five superpop genomes, using Python random seeds from 0 to 14. Then we repeated the experiments using simulated data, using identical setup as described in Sections 2.2 and 2.3. We compared the 15 RandFlow-LD results with vg using a graph built with all variants with $\geq$ 1% allele frequency and showed that the variability of RandFlow-LD is very small compared to the variability in alignment methods in both alignment bias and allelic bias (Figure S10, S11 and S12).

#### Supplementary Figures

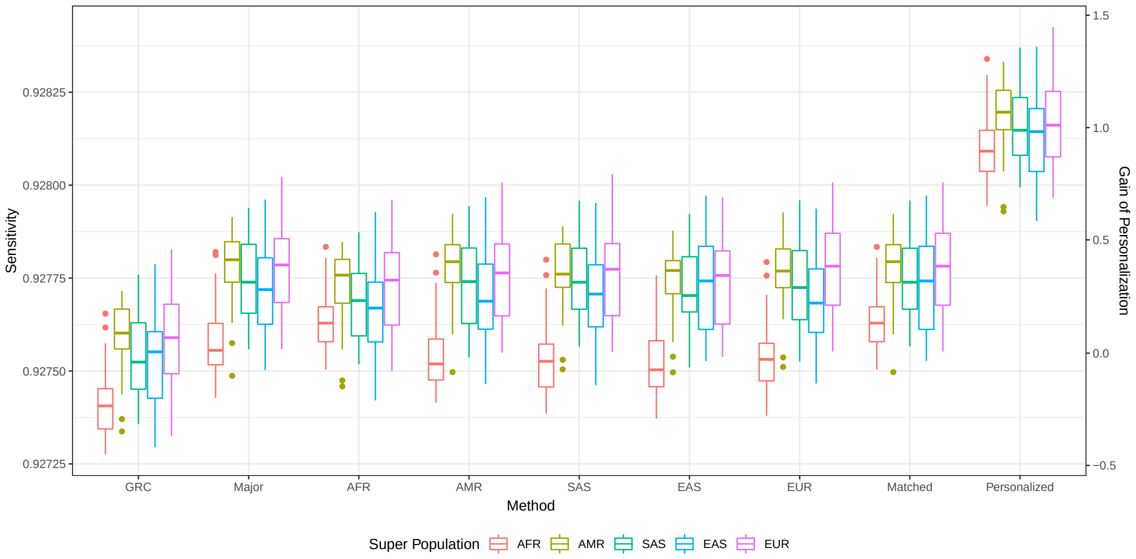


Figure S1: Population-stratified mapping sensitivity using typical methods that all reads are aligned to a single linear reference genome. 2M 100-bp single-end reads are simulated from chromosome 21 and aligned using Bowtie 2. 100 individuals (1M reads per haplotype, both SNVs and indels included) sampled from the 1000 Genomes Project are used for simulation. The *Major* column shows the results using the global major reference. Five superpop major references are labelled with the super population (*AFR*, *AMR*, *SAS*, *EAS* and *EUR*). In the *Matched* column reads are aligned to corresponding ethnicity-matching superpop major reference. The personalized reference genome (*Personalized*) is diploid and others are haploid.

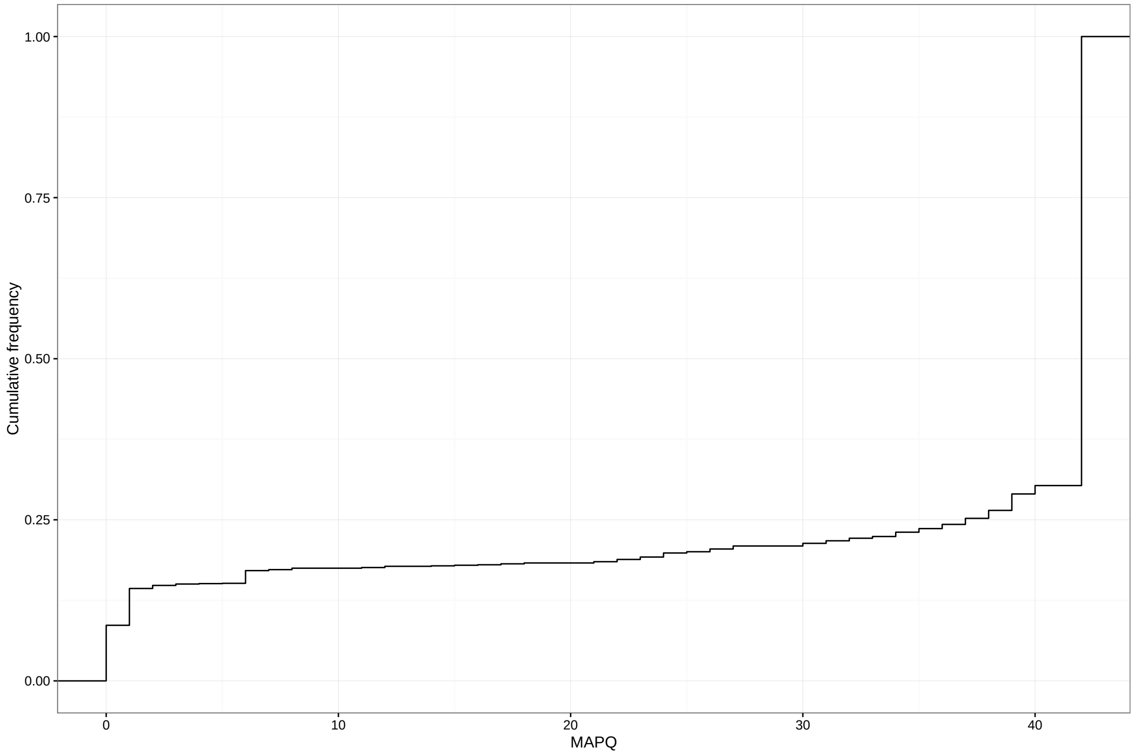


Figure S2: Cumulative frequency of mapping quality (MAPQ) when real whole genome sequencing reads from NA12878 (SRR622457) are aligned to the global major reference genome. More than 82% of reads are aligned with mapping quality $\geq 10$.

We then simulated 2M 100-bp single-end reads using chromosome 21, with NA12878 genotypes, to evaluate alignment correctness. When the MAPQ threshold was set to 10, around 87% of reads were assigned into the committed group and more than 99.98% of these aligned correctly. The alignment sensitivity for the deferred group, where reads are unaligned or aligned with MAPQ lower than 10, was 44.93%.

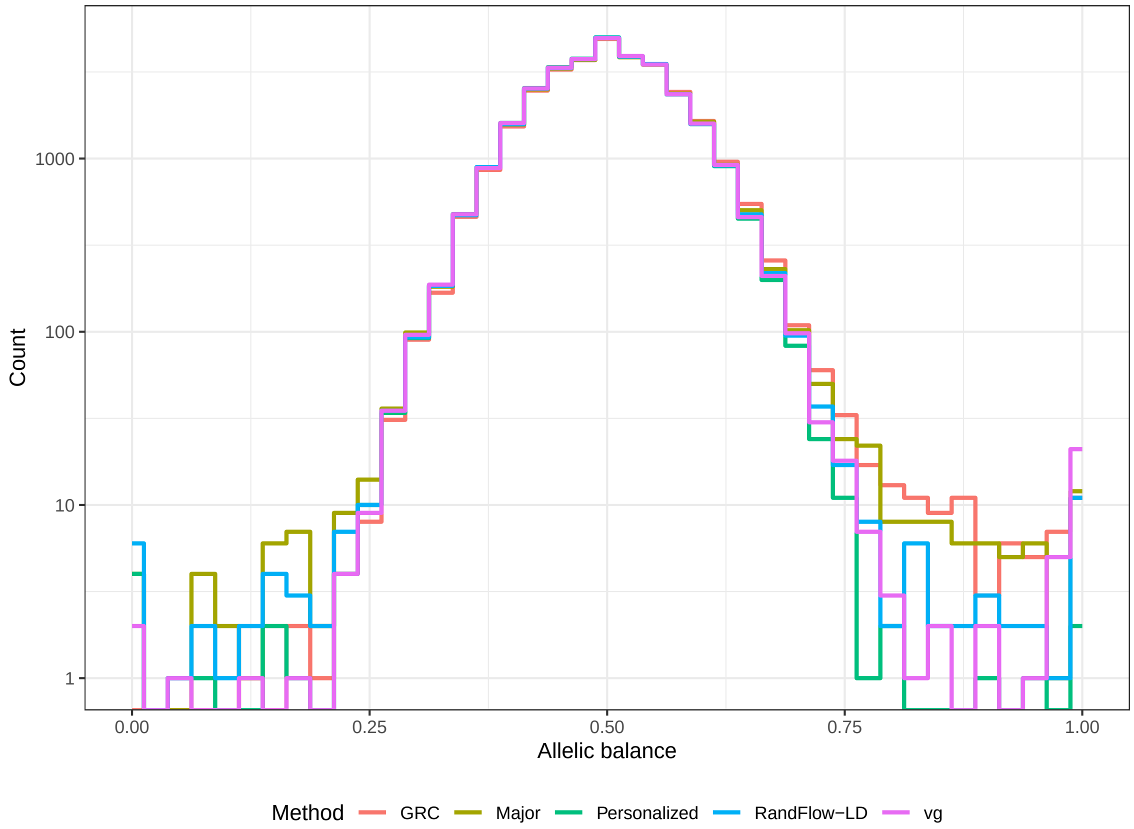


Figure S3: Histograms plotted using 20M 100-bp single-end reads simulated from chromosome 21, GRCh38, with diploid NA12878 genotypes. Chromosome 21 variants with allele frequency $\geq 1\%$ from the 1000 Genomes Project GRCh38 call set are used to build the graph indexes for vg. Numbers of strongly biased sites and REF-to-ALT ratio are shown in Table S4.

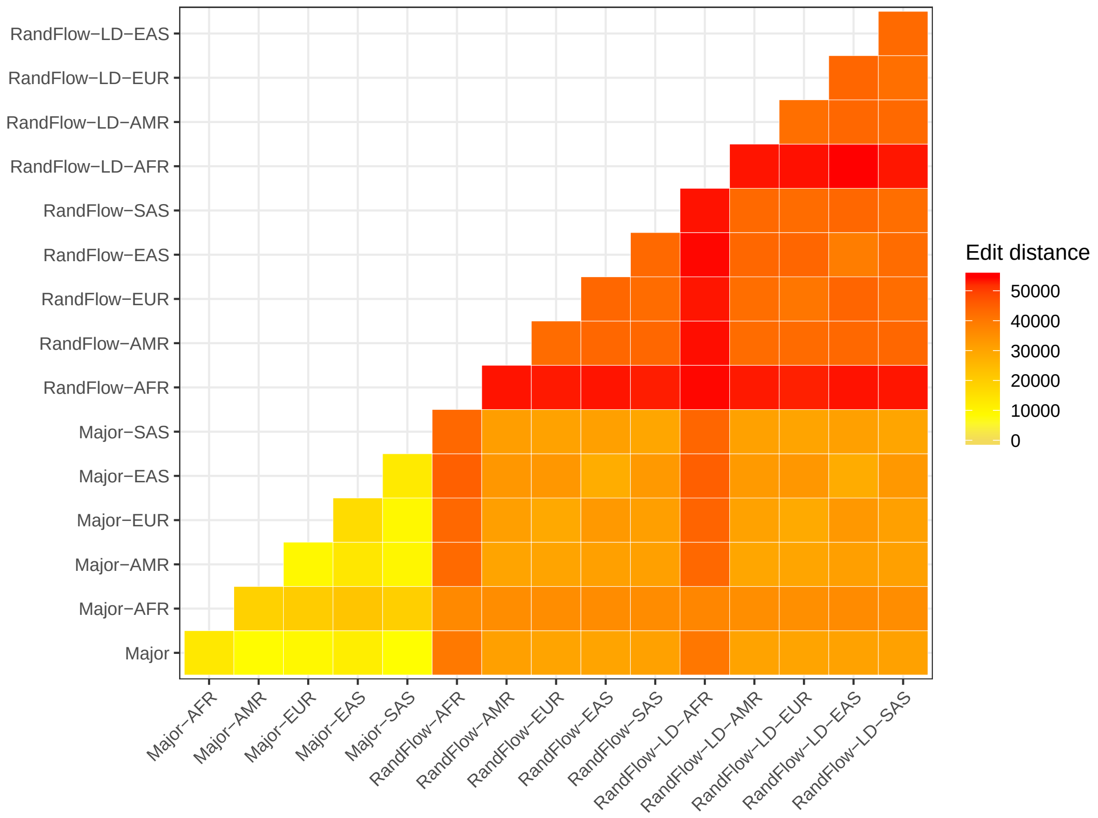


Figure S4: Edit distance, calculated by summing the number of edits for all variants using `bedtools` [45], between global major reference (*Major*), superpop major references (with *Major*- prefixes), stochastic super population genomes (independent-sampling: with *RandFlow*-, phase-preserving with 1000-bp blocks: with *RandFlow-LD*- prefixes).

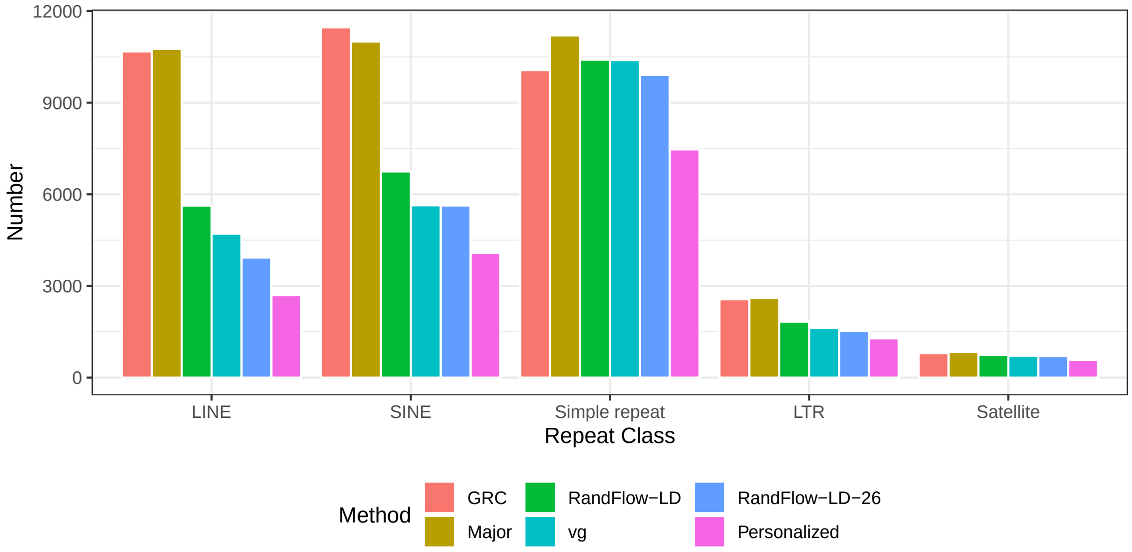


Figure S5: Number of strongly biased HET sites in repetitive elements across the NA12878 genome. Counts are stratified by RepeatMasker class[30]. The reads aligned are the first end of the paired-end reads in SRR622457. Variants with allele frequency $\geq 10\%$ in the 1000 Genomes Project are included in the vg graph.

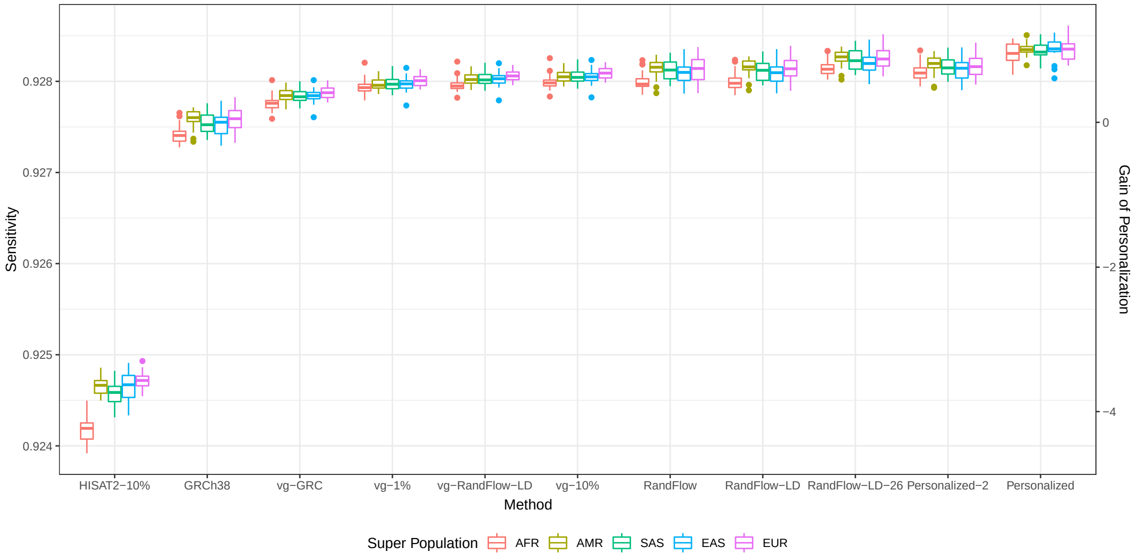


Figure S6: Population-stratified mapping sensitivity including various graph-based approaches. The experiment setup is identical to Figure 2, so as results in columns *GRC*, *RandFlow*, *RandFlow-LD*, *RandFlow-LD-26* and *Personalized*. The *Personalized-2* column shows the results using each personalized haplotype once. The *HISAT-10%* column shows the results using HISAT2 with a graph built using all variants with $\geq$ 10% allele frequency in the 1000 Genomes Project. All columns with prefix ”vg” use aligner vg. In the *vg-GRC* column reads are aligned to linear GRCh38; in the *vg-RandFlow-LD* column reads are aligned to a vg graph built using the RandFlow-LD variant set (Note S1); in the *vg-1%* and *vg-10%* columns reads are aligned to graphs built using variants with allele frequency $\geq$ 1% and 10% respectively. The columns are sorted by median sensitivity.

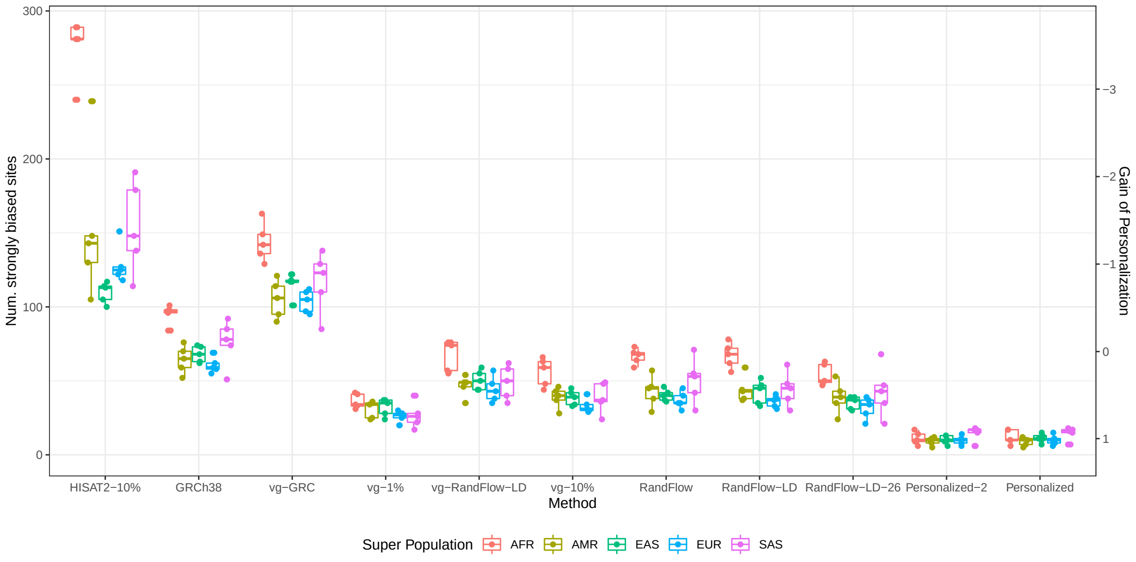


Figure S7: Population-stratified number of strongly biased HET sites including various graph-based approaches. The experiment setup is identical to Figure S6.

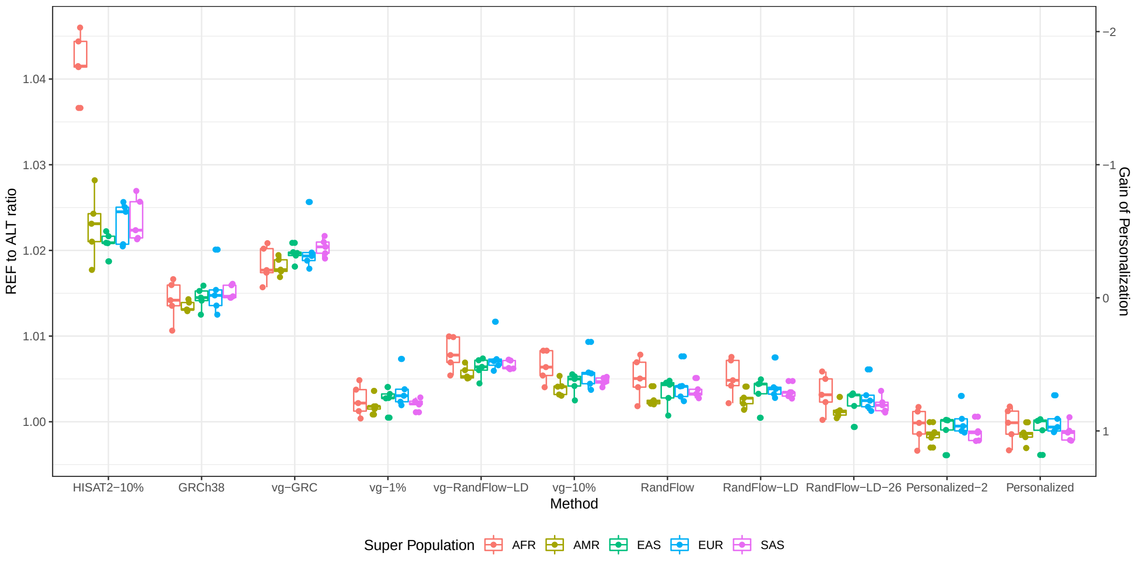


Figure S8: Population-stratified REF-to-ALT ratio including various graph-based approaches. The experiment setup is identical to Figure S6.

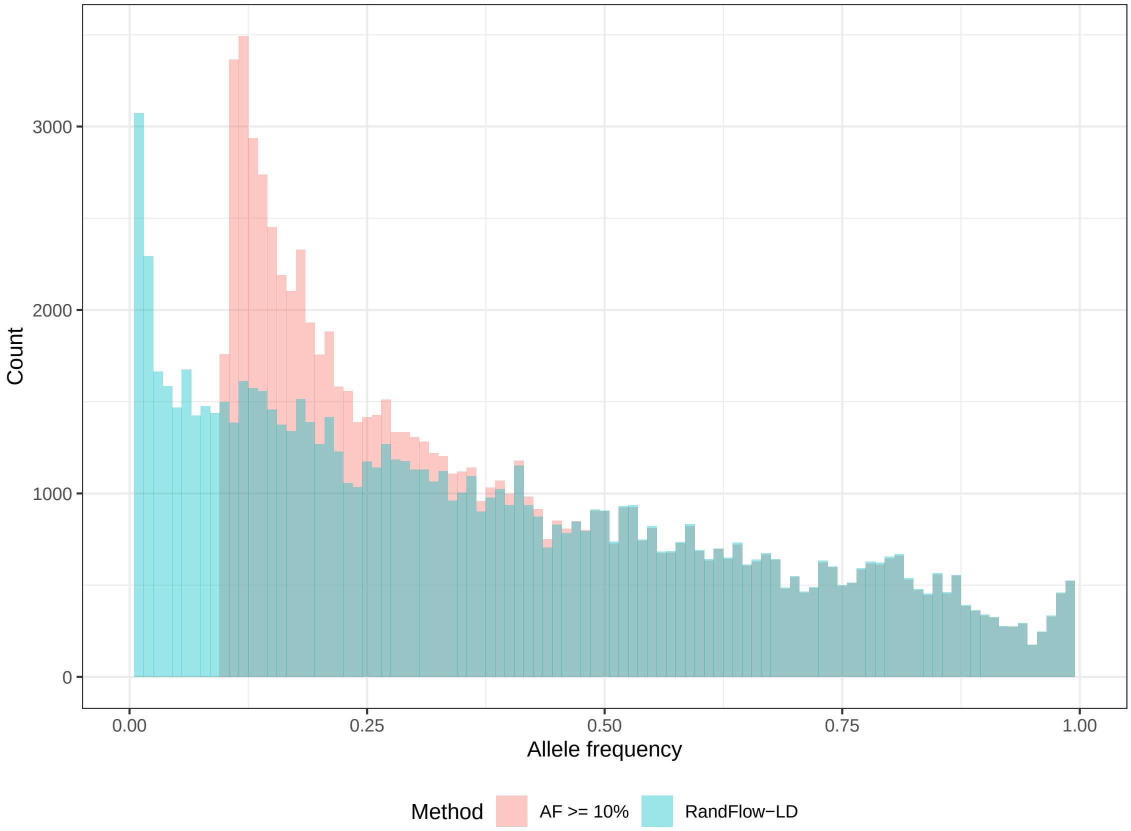


Figure S9: Allele frequency distributions of chromosome 21 RandFlow-LD variants and variants with allele frequency $\geq 10\%$. The RandFlow-LD variant set is the union of major alleles and five superpopulation RandFlow-LD variant sets.

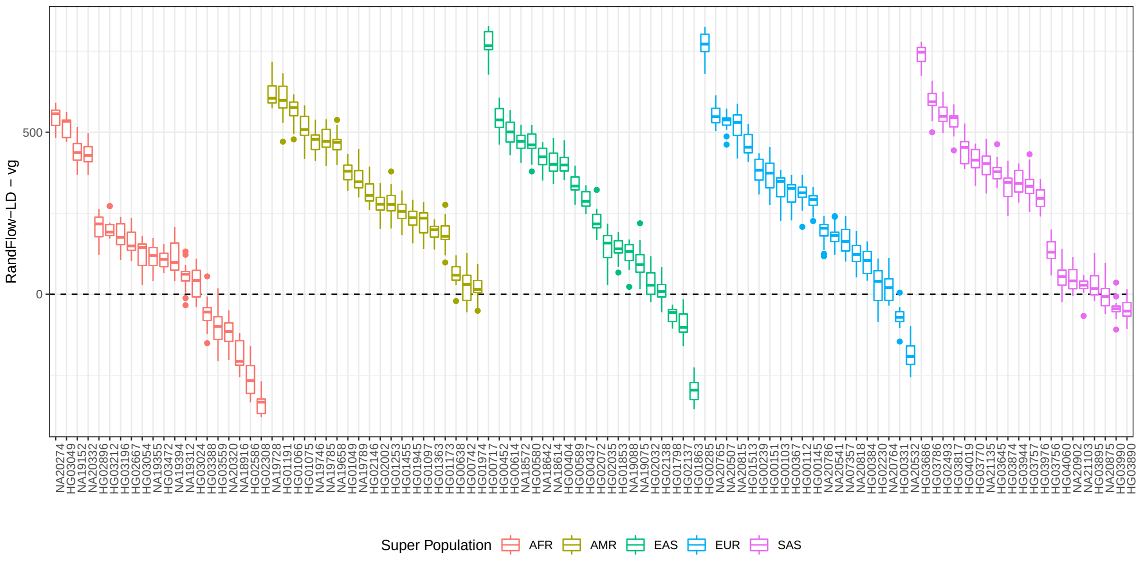


Figure S10: Difference in alignment sensitivity between using vg and 15 sets of RandFlow-LD references, each built with a distinct random seed. The dashed line represents equal sensitivity between two methods. Results within each super population are sorted by median difference in alignment sensitivity. The heights of the boxes convey variability due to the random seed used to select RandFlow genotypes, whereas the vertical spread of same-color boxes conveys variability due to genetic differences between donor individuals. Notably, variability due to random seed is substantially lower compared to variability due to donor-individual genetics.

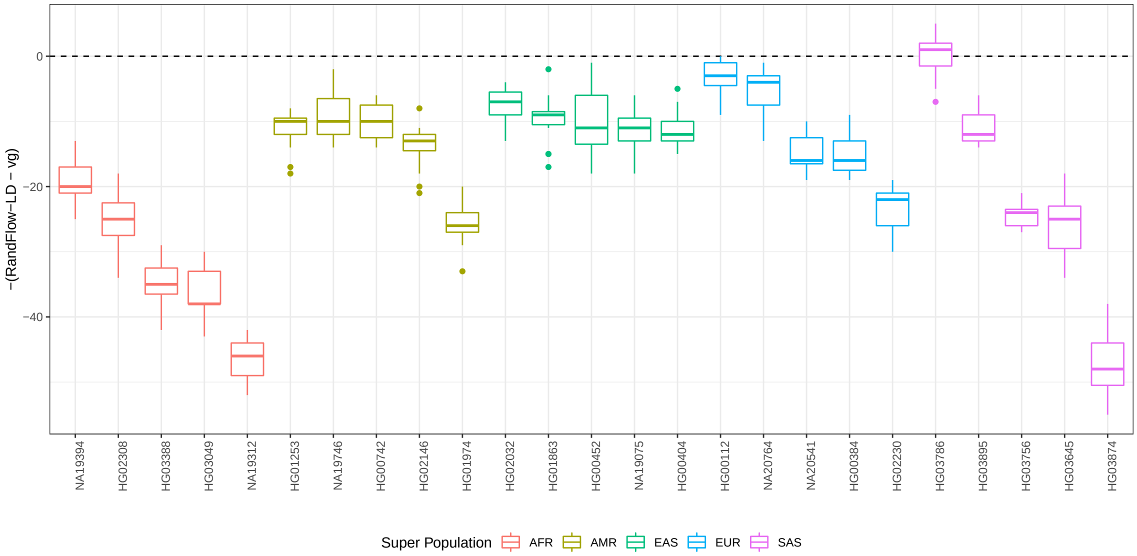


Figure S11: Difference in number of strongly biased sites between using vg and 15 set of RandFlow-LD references, each built with a distinct random seed. The dashed line represents zero difference between two methods. RandFlow-LD outperforms vg if the difference is positive (less number of biased sites). Results within each super population are sorted by median difference in number of strongly biased sites.

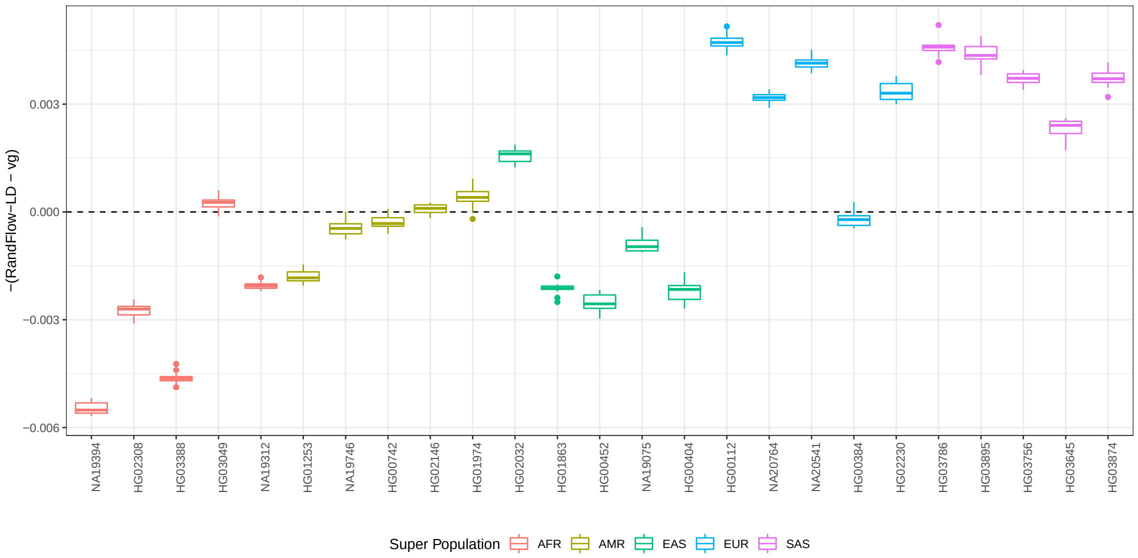


Figure S12: Difference in absolute REF-to-ALT ratio deviation (defined as $|1 - \text{REF-to-ALT}|$) between using vg and 15 sets of RandFlow-LD references, each built with a distinct random seed. The dashed line represents equal REF-to-ALT ratios. RandFlow-LD outperforms vg if the difference is positive. Results within each super population are sorted by median difference in number of strongly biased sites to make it aligned with Figure S11.

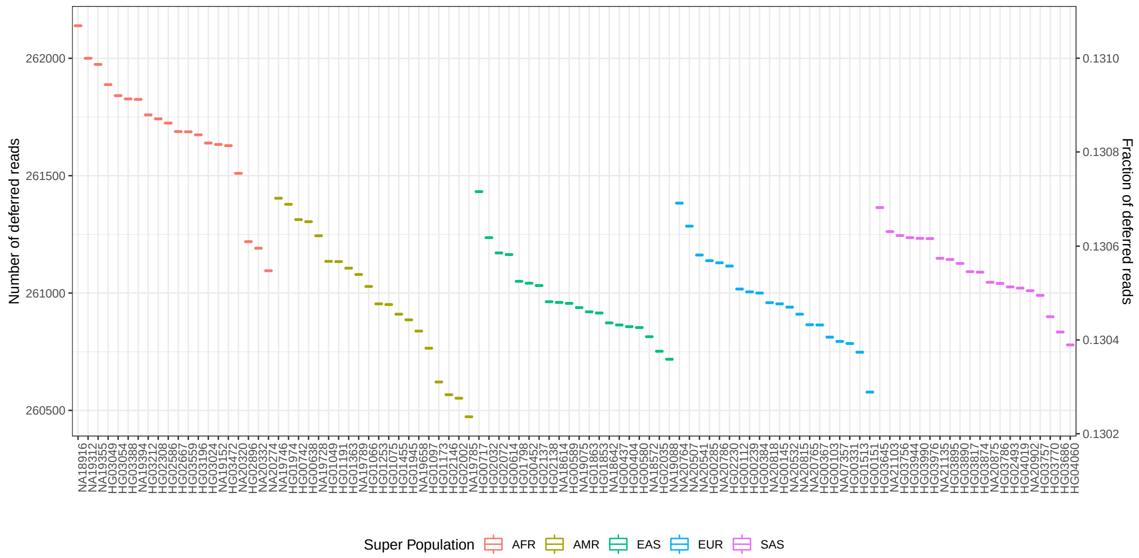


Figure S13: Number of deferred reads (unaligned in the first-pass or aligned with MAPQ $\geq$ 10) when aligning 2M simulated chr21 reads to the global major-allele reference. All reference flow methods discussed in this study — MajorFlow, RandFlow, RandFlow-LD and RandFlow-LD-26 — re-aligned the same set of deferred reads.

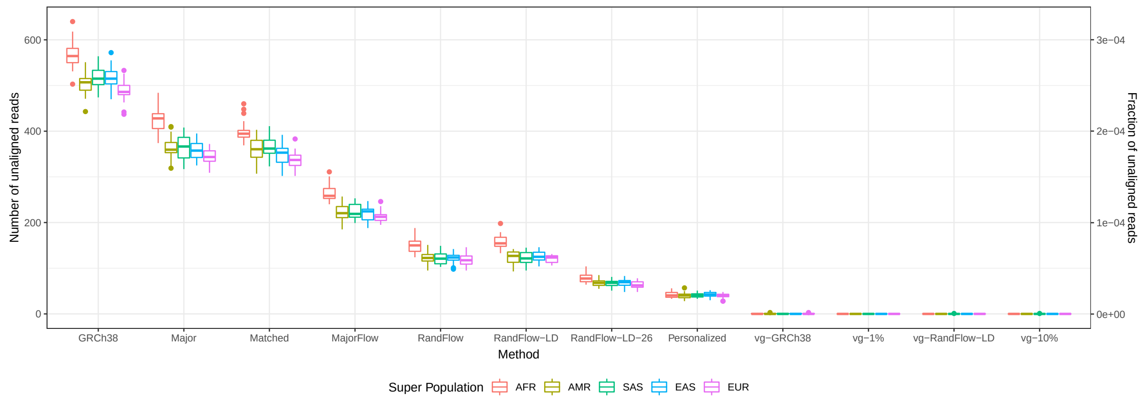


Figure S14: Number of unaligned reads when aligning 2M simulated reads. The experiment setup is identical to Figures 2a and S6.

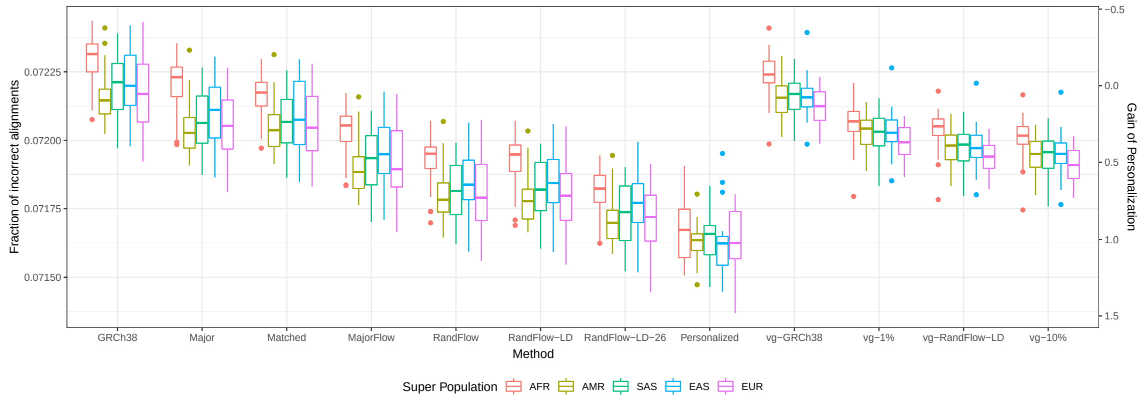


Figure S15: Number of incorrectly aligned reads when aligning 2M simulated reads. The experiment setup is identical to Figures 2a and S6.

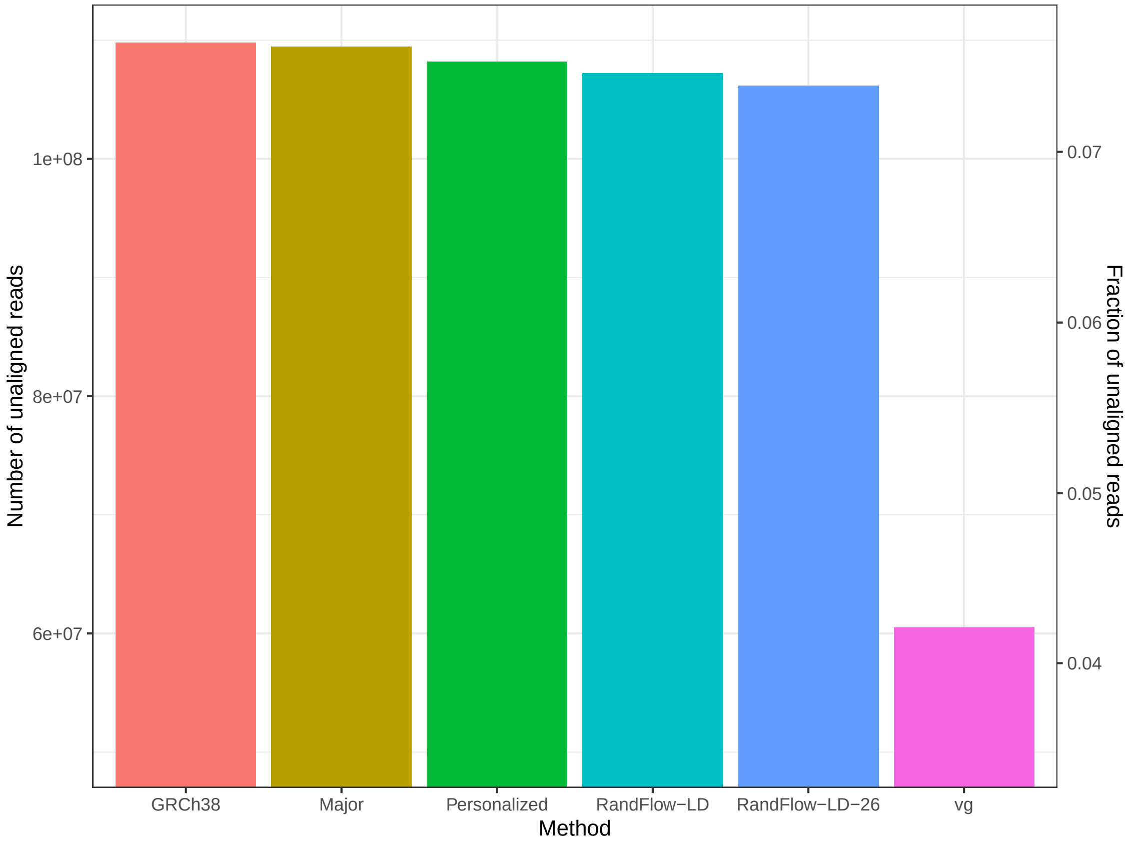


Figure S16: Number of unaligned reads when aligning real reads from SRR622457 to the whole-genome. The experiment setup is identical to Figure 3.

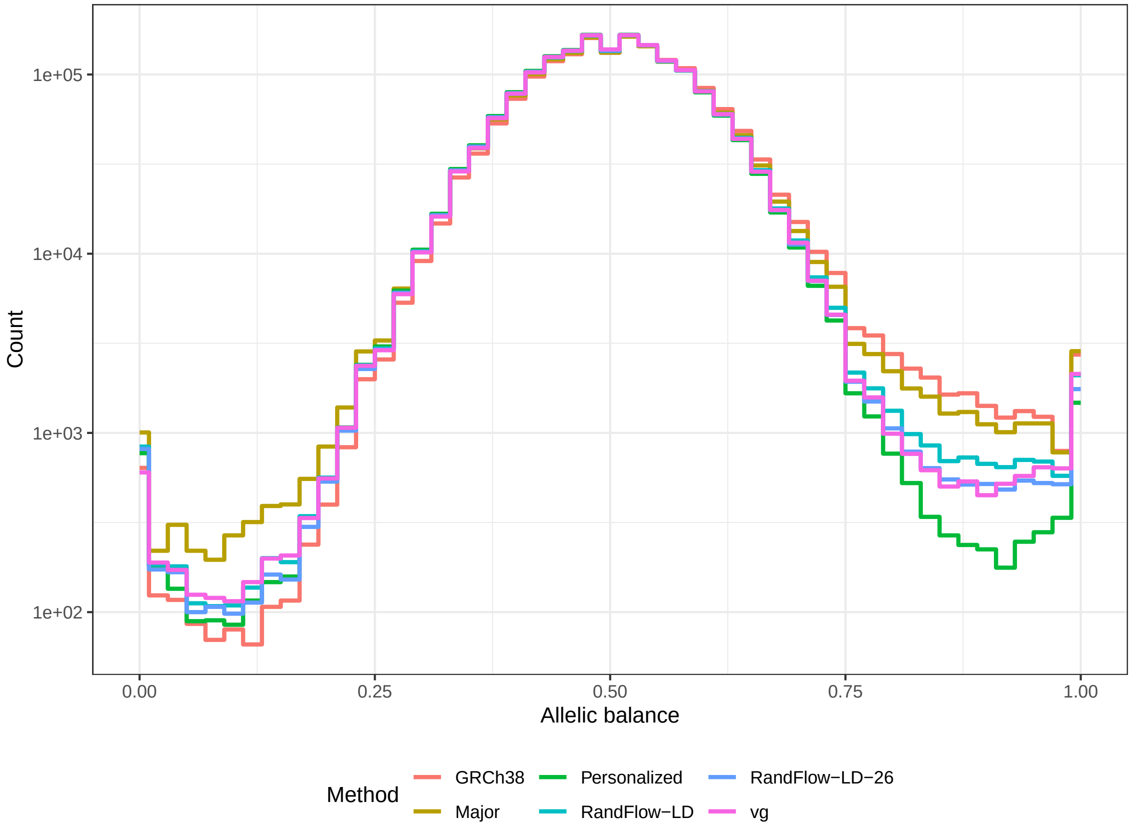


Figure S17: Histograms of allelic balance using a high-coverage real WGS dataset of individual NA12878 (SRR622457) in Genome-in-a-Bottle v3.3.2 high-confidence regions. Experiments are performed using GRCh38 (*GRC*), global major reference (*Major*), diploid personalized genome (*Personalized*), vg using alleles with frequency $\geq 10\%$ (*vg*), reference flow using 1000-bp phased blocks with 5 super populations (*RandFlow-LD*) and reference flow using 1000-bp phased blocks with 26 populations (*RandFlow-LD-26*).

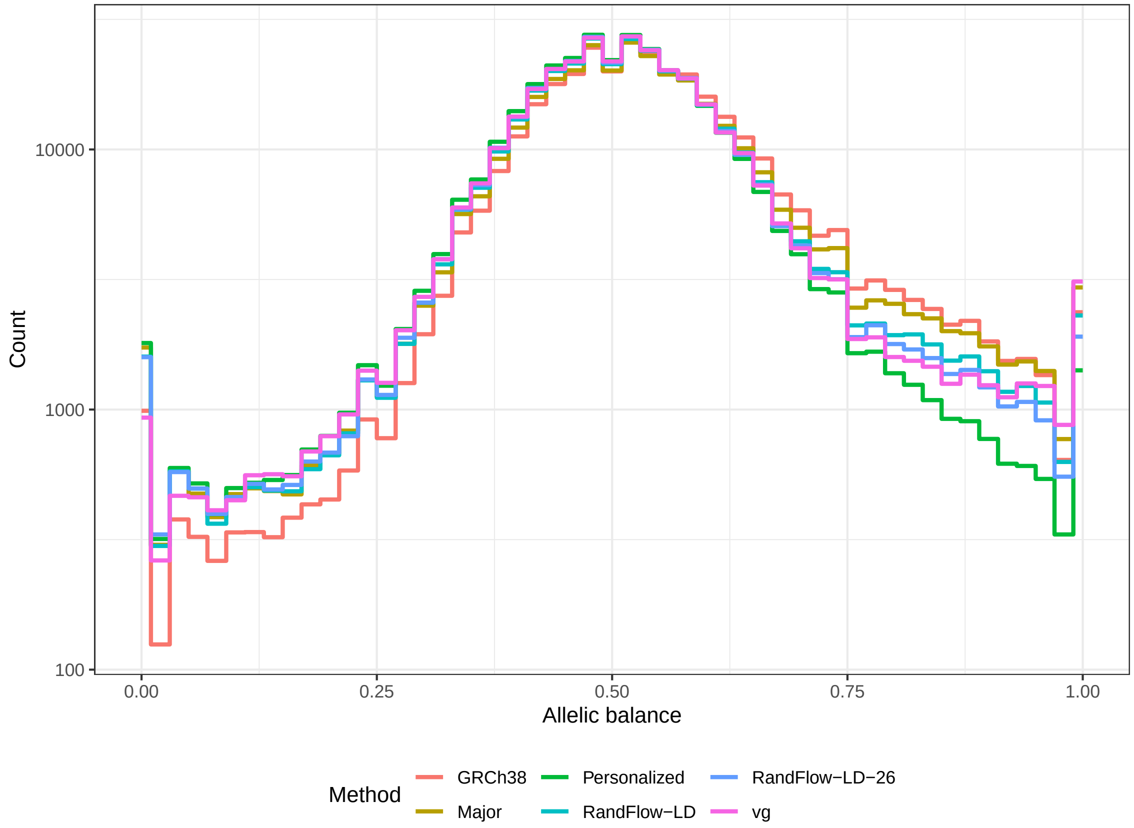


Figure S18: Histograms of allelic balance using a high-coverage real WGS dataset of individual NA12878 (SRR622457) in Genome-in-a-Bottle v3.3.2 low-confidence regions. Experiments are performed using GRCh38 (*GRC*), global major reference (*Major*), diploid personalized genome (*Personalized*), vg using alleles with frequency $\geq$ 10% (*vg*), reference flow using 1000-bp phased blocks with 5 super populations (*RandFlow-LD*) and reference flow using 1000-bp phased blocks with 26 populations (*RandFlow-LD-26*).

#### Supplementary Tables

Table S1: Software versions used.

| Software | Version |
|---|---|
| Bowtie 2[21] | 2.3.4.3 |
| vg[11] | 1.19.0 “Tramutola” |
| Mason2[23] | 2.0.0-beta1 |
| bcftools[31] | 1.9-206-g4694164 |
| samtools[46] | 1.9 |
| Snakemake[42] | 5.5.4 |
| Python | 3.7.3 |
| GNU-time | 1.9 |
| R | 3.6.2 |
| GNU Parallel[47] | 20160622 |

Table S2: A hundred individuals from the 1000 Genomes Project are randomly selected for the simulated experiment. Deeper datasets for allelic bias evaluation are simulated using individuals marked with boldface. Variant statistics for chromosome 21 are reported.

| Sample | Superpopulation | Population | # variants | # SNP | # indel |
|---|---|---|---|---|---|
| **NA19312** | AFR | LWK | 71930 | 63701 | 8229 |
| **NA19394** | AFR | LWK | 73097 | 64709 | 8388 |
| **HG02308** | AFR | ACB | 72263 | 63929 | 8334 |
| **HG03049** | AFR | GWD | 71053 | 62773 | 8280 |
| **HG03388** | AFR | MSL | 71986 | 63652 | 8334 |
| HG03054 | AFR | MSL | 71155 | 62933 | 8222 |
| NA19355 | AFR | LWK | 69661 | 61628 | 8033 |
| HG02667 | AFR | GWD | 71042 | 62896 | 8146 |
| HG03472 | AFR | MSL | 70196 | 62043 | 8153 |
| HG02896 | AFR | GWD | 70096 | 62091 | 8005 |
| NA18916 | AFR | YRI | 70866 | 62619 | 8247 |
| HG03196 | AFR | ESN | 70081 | 62002 | 8079 |
| HG03212 | AFR | MSL | 72923 | 64660 | 8263 |
| NA20320 | AFR | ASW | 66891 | 59114 | 7777 |
| HG02586 | AFR | GWD | 71779 | 63477 | 8302 |
| NA20332 | AFR | ASW | 68034 | 60195 | 7839 |

*Continued on next page*

Table S2 – *Continued from previous page*

| Sample | Superpopulation | Population | # variants | # SNP | # indel |
|---|---|---|---|---|---|
| NA20274 | AFR | ASW | 65038 | 57320 | 7718 |
| NA19152 | AFR | YRI | 72300 | 64003 | 8297 |
| HG03024 | AFR | GWD | 70341 | 62217 | 8124 |
| HG03559 | AFR | MSL | 73777 | 65328 | 8449 |
| **HG01974** | AMR | PEL | 54457 | 47691 | 6766 |
| **HG02146** | AMR | PEL | 56570 | 49596 | 6974 |
| **HG01253** | AMR | CLM | 56730 | 49747 | 6983 |
| **NA19746** | AMR | MXL | 58632 | 51487 | 7145 |
| **HG00742** | AMR | PUR | 57494 | 50477 | 7017 |
| HG00638 | AMR | PUR | 62005 | 54582 | 7423 |
| HG02002 | AMR | PEL | 52990 | 46423 | 6567 |
| NA19789 | AMR | MXL | 58615 | 51378 | 7237 |
| HG01075 | AMR | PUR | 58322 | 51118 | 7204 |
| HG01945 | AMR | PEL | 56900 | 50030 | 6870 |
| HG01097 | AMR | PUR | 59376 | 52151 | 7225 |
| NA19658 | AMR | MXL | 58734 | 51448 | 7286 |
| HG01049 | AMR | PUR | 57367 | 50400 | 6967 |
| HG01455 | AMR | CLM | 56625 | 49676 | 6949 |
| NA19785 | AMR | MXL | 56906 | 49947 | 6959 |
| HG01363 | AMR | CLM | 64792 | 57086 | 7706 |
| HG01191 | AMR | PUR | 57277 | 50262 | 7015 |
| NA19728 | AMR | MXL | 55754 | 48893 | 6861 |
| HG01066 | AMR | PUR | 56643 | 49644 | 6999 |
| HG01173 | AMR | PUR | 57606 | 50577 | 7029 |
| **NA20541** | EUR | TSI | 56993 | 50049 | 6944 |
| **HG00384** | EUR | FIN | 53008 | 46428 | 6580 |
| **HG02230** | EUR | IBS | 54943 | 48284 | 6659 |
| **NA20764** | EUR | TSI | 54921 | 48231 | 6690 |
| **HG00112** | EUR | GBR | 56087 | 49197 | 6890 |
| NA07357 | EUR | CEU | 55650 | 48803 | 6847 |
| HG00285 | EUR | FIN | 53313 | 46605 | 6708 |
| NA20815 | EUR | TSI | 56315 | 49434 | 6881 |
| HG00239 | EUR | GBR | 56735 | 49837 | 6898 |
| HG00151 | EUR | GBR | 54968 | 48142 | 6826 |
| HG00103 | EUR | GBR | 55580 | 48743 | 6837 |
| NA20786 | EUR | TSI | 59886 | 52673 | 7213 |
| NA20507 | EUR | TSI | 57635 | 50660 | 6975 |
| NA20532 | EUR | TSI | 55229 | 48348 | 6881 |

*Continued on next page*

Table S2 – *Continued from previous page*

| Sample | Superpopulation | Population | # variants | # SNP | # indel |
|---|---|---|---|---|---|
| HG00331 | EUR | FIN | 54977 | 48254 | 6723 |
| NA20818 | EUR | TSI | 57044 | 50007 | 7037 |
| HG00145 | EUR | GBR | 56072 | 49173 | 6899 |
| HG00367 | EUR | FIN | 57514 | 50442 | 7072 |
| HG01513 | EUR | IBS | 55368 | 48614 | 6754 |
| NA20765 | EUR | TSI | 58726 | 51531 | 7195 |
| **HG01863** | EAS | KHV | 55137 | 48360 | 6777 |
| **HG00404** | EAS | CHS | 55263 | 48390 | 6873 |
| **NA19075** | EAS | JPT | 56286 | 49379 | 6907 |
| **HG02032** | EAS | KHV | 55801 | 48905 | 6896 |
| **HG00452** | EAS | CHS | 57862 | 50877 | 6985 |
| HG01798 | EAS | CDX | 55114 | 48351 | 6763 |
| NA18572 | EAS | CHB | 55384 | 48552 | 6832 |
| HG02137 | EAS | KHV | 56923 | 50006 | 6917 |
| HG00614 | EAS | CHS | 55597 | 48708 | 6889 |
| HG00589 | EAS | CHS | 56125 | 49265 | 6860 |
| HG00437 | EAS | CHS | 57259 | 50351 | 6908 |
| HG02072 | EAS | KHV | 58807 | 51691 | 7116 |
| NA19088 | EAS | JPT | 56111 | 49149 | 6962 |
| HG01853 | EAS | KHV | 55178 | 48333 | 6845 |
| NA18614 | EAS | CHB | 56190 | 49212 | 6978 |
| HG02138 | EAS | KHV | 56748 | 49798 | 6950 |
| HG00580 | EAS | CHS | 57456 | 50388 | 7068 |
| NA18642 | EAS | CHB | 55687 | 48833 | 6854 |
| HG00717 | EAS | CHS | 55592 | 48757 | 6835 |
| HG02035 | EAS | KHV | 56565 | 49596 | 6969 |
| **HG03756** | SAS | STU | 58530 | 51352 | 7178 |
| **HG03874** | SAS | ITU | 58693 | 51625 | 7068 |
| **HG03645** | SAS | STU | 58696 | 51612 | 7084 |
| **HG03895** | SAS | STU | 50966 | 44682 | 6284 |
| **HG03786** | SAS | ITU | 58229 | 51133 | 7096 |
| HG03817 | SAS | BEB | 56287 | 49307 | 6980 |
| NA20902 | SAS | GIH | 57512 | 50511 | 7001 |
| HG04060 | SAS | ITU | 56514 | 49641 | 6873 |
| HG04019 | SAS | ITU | 57875 | 50819 | 7056 |
| HG03770 | SAS | ITU | 59557 | 52338 | 7219 |
| HG03990 | SAS | STU | 59801 | 52622 | 7179 |
| HG03686 | SAS | STU | 57325 | 50370 | 6955 |

*Continued on next page*

Table S2 – *Continued from previous page*

| **Sample** | **Superpopulation** | **Population** | **# variants** | **# SNP** | **# indel** |
|---|---|---|---|---|---|
| HG03757 | SAS | STU | 60150 | 52872 | 7278 |
| NA21103 | SAS | GIH | 57945 | 50894 | 7051 |
| HG03976 | SAS | ITU | 56991 | 49993 | 6998 |
| NA21135 | SAS | GIH | 57912 | 50857 | 7055 |
| HG02493 | SAS | PJL | 58844 | 51664 | 7180 |
| HG03890 | SAS | STU | 58140 | 50998 | 7142 |
| HG03944 | SAS | STU | 58349 | 51186 | 7163 |
| NA20875 | SAS | GIH | 58039 | 51000 | 7039 |

Table S3: Variant statistics of NA12878 on chromosome 21

| Sample | Superpopulation | Population | # variants | # SNP | # indel |
|---|---|---|---|---|---|
| NA12878 | EUR | CEU | 56905 | 50012 | 6893 |

Table S4: Allelic balance of the simulated NA12878 chr21 dataset. 20 million 100-bp single-end reads are aligned using different methods.

| Method | # REF biases | # ALT biases | Total | REF to ALT ratio |
|---|---|---|---|---|
| GRCh38 | 73 | 4 | 77 | 1.0141 |
| Major | 58 | 25 | 83 | 1.0064 |
| MajorFlow | 54 | 22 | 76 | 1.0053 |
| RandFlow-LD | 29 | 20 | 49 | 1.0027 |
| vg | 33 | 5 | 38 | 1.0031 |
| Personalized | 4 | 9 | 13 | 0.9991 |

Table S5: Numbers of variants across the whole-genome in augmented references. The RandFlow-LD set is the union of *Major* and five superpopulation RandFlow-LD variant sets.

| Population | Number of variants |
|---|---|
| Personalized (diploid) | 3,903,552 |
| Major | 1,998,961 |
| RandFlow-LD | 6,471,383 |
| AFR (RandFlow-LD) | 3,155,143 |
| AMR (RandFlow-LD) | 2,769,584 |
| EAS (RandFlow-LD) | 2,786,606 |
| EUR (RandFlow-LD) | 2,711,876 |
| SAS (RandFlow-LD) | 2,763,946 |
| $\geq$10% allele frequency | 6,267,736 |
| $\geq$1% allele frequency | 13,511,758 |
